# Genomic detection of domoic acid biosynthesis uncovers expanded toxigenic diversity across *Pseudo-nitzschia* diatoms

**DOI:** 10.64898/2026.10.23.761457

**Authors:** Steffaney M. Wood-Rocca, Zoltán Füssy, Monica Thukral, Andres Mauricio Caraballo-Rodriguez, Hong Zheng, Shelly A. Wanamaker, Jennifer M. Polinski, Alexandra DeSmidt, Nathalie Choisne, Brian D. Bill, Hannah McGrath, Freya Hammar, Florian Maumus, Katherine A. Hubbard, Pieter C. Dorrestein, Tom O. Delmont, Matthew J. Harke, Bradley S. Moore, Andrew E. Allen

## Abstract

Domoic acid (DA) production by diatom genus *Pseudo-nitzschia* causes recurrent harmful algal blooms with major consequences for marine ecosystems, fisheries, and human health. Predicting DA risk remains challenging because toxin production varies across morphologically similar and cryptic lineages. Consequently, morphology or single marker genes often fail to resolve toxigenic potential. Here, we expand genomic resources for *Pseudo-nitzschia* by generating and compiling high-quality genomes spanning diverse species and biogeographic origins, establishing a genus-wide phylogenomic framework to investigate the distribution and evolution of the DA biosynthesis (*dab*) gene cluster. Among chemically characterized strains, the *dab* pathway was detected in species only with confirmed DA production. While the cluster exhibits a conserved core structure, several lineages display signatures of ongoing evolution, including intronization, locus rearrangement, and pseudogenization. One striking example of *dab* variation occurs in toxigenic *P. australis* strains, in which its biochemically validated isodomoic acid A synthase-encoding gene *dabC* is located outside the *dab* locus. Metatranscriptomic surveys reveal widespread environmental *dab* expression across size classes and species, implicating taxa not typically associated with environmental DA events, such as *P*. cf. *obtusa* and *P*. cf. *plurisecta*. To connect *dab* expression with chemical output, nutrient limitation experiments in the historically inconsistent DA producer *P. pseudodelicatissima* combined transcriptional profiling and nontargeted metabolomics demonstrating coordinated *dab* expression across distinct co-expression modules and DA accumulation in response to silicon limitation. Together, these results refine the phylogenetic scope of DA production and provide a genome-resolved basis for improved molecular monitoring of toxigenic diversity.

## Introduction

*Pseudo-nitzschia* is a cosmopolitan diatom genus that forms harmful algal blooms with far-reaching economic, ecological, and human health consequences. Of the more than 50 described species, roughly half are capable of producing the neurotoxin domoic acid (DA) (1). DA is the causative agent of amnesic shellfish poisoning in humans and responsible for extensive fisheries closures, marine animal poisonings, and significant economic losses worldwide (2).

Despite decades of research, predicting DA production remains a challenge because uncertainty arises from multiple biological scales. First, several *Pseudo-nitzschia* species are considered inconsistent DA producers because toxin production is only observed in some isolates (1) or under certain environmental conditions (3–7). Second, many species belong to cryptic or pseudo-cryptic complexes that cannot be distinguished morphologically or using single genetic markers (8, 9). Third, DA production has traditionally been associated with the large-celled seriata size class, whereas small-celled members of the delicatissima size class have been considered low risk or non-toxigenic (8, 9). Together, these factors complicate efforts to infer toxigenic potential from species identity alone.

The discovery of the DA biosynthesis (*dab*) gene cluster provided a molecular basis for predicting DA production (10, 11). The *dab* cluster contains four genes encoding N-prenyltransferase DabA, unknown-function protein DabB, kainoid synthase DabC, and cytochrome P450 DabD. In *P. multiseries*, DabACD have been demonstrated to collectively convert geranyl diphosphate and L-glutamate to isodomoic acid A en route to DA (10). This molecular framework challenged the long-held view, established by early physiological studies, that environmental conditions alone determine whether a *Pseudo-nitzschia* strain produces DA (1, 2, 6, 12).

Recent studies have increasingly established the presence or absence of the *dab* gene cluster across *Pseudo-nitzschia* species. While initially identified in high DA producers, the *dab* cluster has also been found in an inconsistent DA-producing species *P. cuspidata* (13) and in low DA-producing strains of *P. multistriata* (7). The conserved gene synteny facilitates detection of *dab* genes in environmental genomic and transcriptomic datasets (14–16). Importantly, sequence variation in the *dab* genes, particularly kainoid synthase *dabC* genes, encodes distinct DA chemotypes, as validated by biochemical assays (17, 18). This enables DA chemistry to be predicted from gene sequence alone, establishing a direct link between genomic potential and toxin structure (11, 19). As species distributions shift with global change, inferring toxigenicity directly from environmental DNA and RNA datasets becomes increasingly critical (20, 21).

Despite these advances, predicting environmental DA production remains challenging due to genomic insufficient genomic reference coverage of *Pseudo-nitzschia* diversity, and the variability of pathway activation and toxin accumulation with physiological state. Single-marker approaches lack the phylogenomic resolution needed to interpret *dab* gene function, evolution, and distribution across species. As a result, current monitoring frameworks struggle to link species identity, genetic toxin potential, and realized toxin production. While field metatranscriptomic surveys (15) and targeted studies (22) have begun to uncover *dab* cluster diversity and expression patterns, including observations that *dabA* and silicon transporter *sit1* co-expression precede DA toxicity in a *P. australis* bloom (11), a comprehensive understanding of dab-containing lineages, pathway architecture, and physiological context is still needed to interpret and extend these molecular signals across the diverse *Pseudo-nitzschia*.

Herein, we integrate comparative genomics, environmental transcriptomics, and culture-based experiments to disentangle variability in toxin production, cryptic diversity, and environmental regulation of the DA biosynthesis pathway in *Pseudo-nitzschia*. We expand genomic resources by generating high-quality genomes from a diverse array of species and geographic isolates. Enabling a comprehensive, genus-wide assessment of *dab* gene distribution to resolve species-specific patterns of *dab* gene expression across global metatranscriptomes. Additionally, by conducting nutrient-limitation experiments with the environmentally abundant and historically inconsistent DA producer *P. pseudodelicatissima*, we elucidate the temporal relationship between *dab* gene expression, DA biosynthetic intermediates, and DA accumulation. Collectively, our integrative approach links species identity, genomic potential, environmental activity, and realized DA production, providing a robust foundation for interpreting toxin dynamics and improving molecular monitoring of toxigenic diversity in *Pseudo-nitzschia*.

## Results

### Towards a comprehensive genomic resource for toxigenic *Pseudo-nitzschia*

To date, genomes available for *Pseudo-nitzschia* have not comprehensively covered all species suspected of DA production. To address this knowledge gap, we isolated cell lines from various locations and acquired additional material from culture collections for genome sequencing and assembly (Fig. S1A, Table S1). For each of these cultures we obtained scaffold- and contig-level assemblies with high completeness according to conserved ortholog scores (Fig.1, Dataset S1-2). We complemented our newly generated data with additional available genomic, metagenome-assembled genomic (MAG, 23), and transcriptomic data from various sources (Dataset S2), and used this dataset for downstream phylogenomic and comparative analyses.

**Figure 1.**
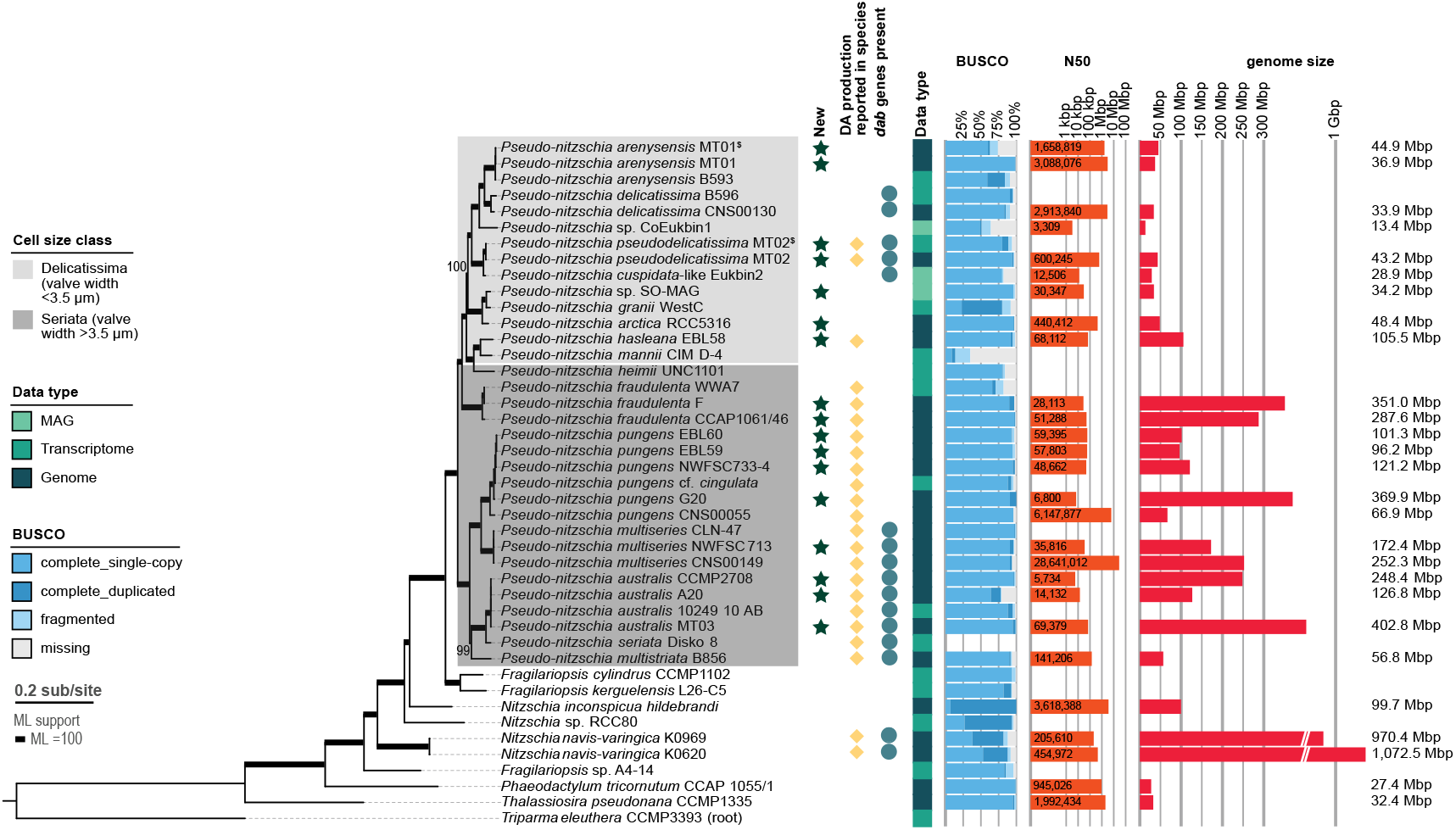
The toxigenic *Pseudo-nitzschia* genomic resource. An IQ-TREE maximum-likelihood phylogenomic tree from 217 genes / 87,978 positions, with ultra-fast bootstrap support (full support shown as thick nodes). Tree rooted with *Triparma eleuthera*. Annotations mark if a taxon is a new dataset, a domoic acid (DA) producer, and the type of resource (genomic, metagenomic, or transcriptomic, see legend). Additional columns describe resource completeness (% BUSCOs), contiguity (N50), and size (for genomes). ^$^ – alternative pilot data for comparison. Interactive version of the figure available on iTOL (https://itol.embl.de/tree/107207231143134911765429885).

Genome sizes varied considerably (34-403 Mbp), with species of the smaller-celled delicatissima size class generally harboring the smallest genomes (34-45 Mbp). Most of the larger genomes, in comparison, exhibited expanded repetitive content (Dataset S2). Analyses of protein family content and evolution found little evidence for significant expansion or contraction of protein familes in the common ancestor of *Pseudo-nitzschia* or in the major clades (Fig. S2). The core *Pseudo-nitzschia* proteome is estimated to comprise 9,874 genes with a Pfam annotation, based on the average number across the 16 analyzed species (excluding repeat-related families; Dataset S3). Similarly, OrthoFinder estimates an average of 11,323 genes in core *Pseudo-nitzschia* orthogroups, defined as those present in more than 75% of species. The largest gene families are associated with signal transduction, transcriptional regulation, protein turnover, and metabolite transport/biosynthesis (Dataset S3). These findings suggest that evolutionary diversification in *Pseudo-nitzschia* was driven more by changes in transcriptional regulation than by expansion of specific gene families.

The search for conserved phylogenetic markers yielded 217 proteins, which concatenated to 87,978 positions. The phylogenomic tree fully converged and resolved the relationships between *Pseudo-nitzschia* species. A deep split was identified between the *P. seriata* complex (encompassing *P. pungens, P. multiseries, P. australis, P. seriata*, and *P. multistriata*) and a second clade, which included *P. fraudulenta, P. granii, P. heimii* and historically recognized species of the *P. delicatissima*/*pseudodelicatissima* complexes (*P. arenysensis, P. delicatissima, P. cuspidata, P. pseudodelicatissima, P. arctica, P. hasleana, P. mannii*; (Fig. S1B; 24). The relationship between *Pseudo-nitzschia* spp., *Fragilariopsis* spp., *Nitzschia* spp., and other outgroups in the analysis was also fully resolved. Importantly, the phylogenomic tree was highly congruent with the phylogeny based on ribosomal DNA operons (spanning the region roughly between 1510R (25) and 28SR (11) primer binding sites, 1,678 positions; Fig. S1A, Supplementary Data). The latter provided (sub)species-level resolution in the absence of genomic data from lineages not presently in culture (Fig. S1A). In the past, structure-based ITS2 data provided (sub)species resolution but insufficient support for deep nodes within the genus, whereas V4/V9 sequences were found insufficient for either species or backbone resolution (26).

Our phylogenomic analysis encompassed several new lineages harboring *dab* gene clusters, particularly *P. pseudodelicatissima* and closely related *Pseudo-nitzschia* sp. EukBin2 (hereafter termed *P. cuspidata*-like EukBin2 based on phylogenetic position), and *P. seriata* Disko8. This finding allowed us to explore patterns of the *dab* gene cluster distribution in greater depth than previously possible.

### Domoic acid biosynthesis gene cluster across *Pseudo-nitzschia* species

Comparative analysis of *dab* gene cluster architecture across *Pseudo-nitzschia* lineages revealed that biosynthetic capacity extends beyond the traditional size-class paradigm and, among the examined strains, was restricted to those that produced DA (Fig. 2A; Fig. S3).

**Figure 2.**
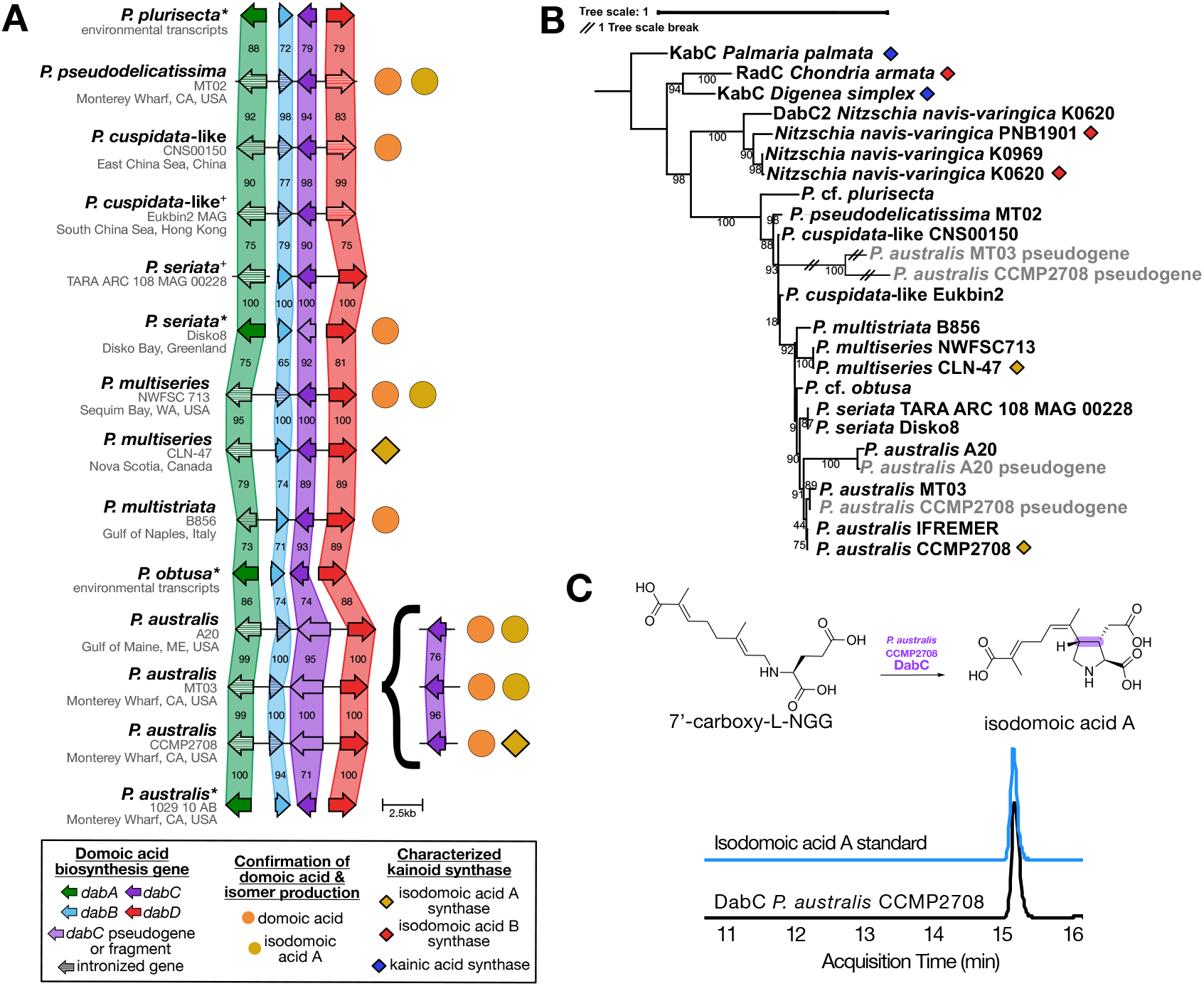
Structural variation and pseudogenization within the *dab* biosynthetic gene cluster across *Pseudo-nitzschia* isolates. **(A)** Synteny comparison of the domoic acid biosynthetic gene cluster (*dabA–D*) across *Pseudo-nitzschia* genomes, metagenome-assembled genomes (indicated by +), and transcriptomes (indicated by *) from diverse geographic regions. The *P. seriata* TARA ARC MAG cluster is interrupted between *dabA* and *dabB* as indicated by non-contiguity. Genes are shown as colored arrows indicating transcriptional orientation, with percent amino acid identity to the neighboring sequence indicated within each gene block. Symbols to the right denote domoic acid and isomer production, and kainoid synthase status. Insets to the right of *P. australis* highlight examples of truncated or disrupted kainoid synthase *dabC* genes and intact copies outside the gene cluster. **(B)** Maximum likelihood phylogeny of the kainoid synthase genes, including intact copies and pseudogenes, and gene losses. **(C)** Extracted ion chromatogram of isodomoic acid A for enzyme activity assay of non-clustered *P. australis* CCMP2708 DabC with 7’-carboxy-L-NGG, compared with isodomoic acid A standard.

*Pseudo-nitzschia* species have historically been divided into the large “seriata” size class (i.e., *P. seriata* complex, *P. fraudulenta, P. heimii*, and *P. subpacifica*) with frustule widths >3 μm and the small “delicatissima” (<3 μm) size class (24). The seriata class is closely associated with DA production and targeted in monitoring programs (27–29). In contrast, the delicatissima class generally produces no or lower cellular DA quotas, although dense blooms can reach comparable toxicity (2). Several members of the delicatissima size class, including *P. pseudodelicatissima* and *P. cuspidata*, phylogenetically belong to cryptic or pseudo-cryptic species complexes and have long been regarded as inconsistent DA producers. Among inconsistent DA producers examined here, only strains of *P. pseudodelicatissima* (n=1) and *P. cuspidata* (n=1) produced isodomoic acid A and DA and harbored the *dab* cluster, whereas strains of *P. fraudulenta* (n=2) and *P. pungens* (n=5) did not.

The *dab* cluster is conserved in gene composition, order, and orientation, with *dabA* and *dabC* oriented in one direction and *dabB* and *dabD* in the opposite direction. Despite the geographic difference among conspecific strains, the Dab protein sequences display between 95 to 100 percent amino acid identity, particularly in established high DA-producing species *P. australis, P. multiseries*, and *P. seriata*. Conservation remains high (90-100 percent identity) in inconsistent DA producers, such as *P. cuspidata*-like and *P. pseudodelicatissima* (Fig. 2A). This broader range may reflect taxonomic ambiguity within this group. Alignment of the genomic sequences with both sequencing reads and assembled transcripts identified putative introns in *dabB* and *P. cuspidata*-like/*pseudodelicatissima* clade-specific introns in *dabD* (Fig. 2A, Fig. S4). Previously, only *dabA* was recognized to contain an intron (5).

Individual *dab* gene phylogenies largely mirrored the genome-resolved phylogeny, including separation of the size classes (Fig. S5-6; Fig. 2B). Taken together, phylogenetic and genomic evidence suggest a high prevalence of DA production in an entire lineage of smaller size class *Pseudo-nitzschia*; henceforth, we refer to the species of this lineage as the *pseudodelicatissima*-like *dab* clade.

Except for *P. australis, dabA, dabB, dabC*, and *dabD* are consistently co-localized within a single locus in toxigenic lineages. Contrastingly, in all three *P. australis* strains, a *dabC* pseudogene is co-localized with the cluster, while the intact, full-length *dabC* open reading frame resides elsewhere in the genome. Some strains have an additional *dabC* pseudogene, also found outside the cluster, with a non-sense mutation in the coding sequence. Notably, *P. australis* pseudogenes in strains MT03 and CCMP2708 are the most divergent and fall within the *P. cuspidata*-like clade as opposed to the *P. australis* clade (Fig. 2B). Heterologous expression and *in vitro* enzyme assays verified the function of the *P. australis* CCMP2708 non-clustered *dabC* as an isodomoic acid A synthase (Fig. 2C, Fig. S7).

Phylogenetic analysis of kainoid synthase sequences, including stop codon-containing *dabC* pseudogenes, results in a cohesive *Pseudo-nitzschia* DabC clade (Fig. 2B, S6B), distinct from isodomoic acid B and kainic acid synthase enzymes found in red algae and in the benthic diatom *N. navis-varingica* (10, 17, 18, 30). Therefore, all *Pseudo-nitzschia* kainoid synthase enzymes are inferred to be isodomoic acid A synthases, which aligns with the detection of isodomoic acid A and domoic acid in culture extracts (Fig. S3). While isodomoic acid A and domoic acid production could not be confirmed from strains of publicly available assemblies analyzed here, production of at least isodomoic acid A can be inferred from the kainoid synthase phylogeny (Fig. S6B).

### Environmental expression reveals domoic acid production potential in cryptic lineages

Our MAG- and metatranscriptome (metaT)-based comparisons show that environmental data harbor considerable, underexploited potential for novel *dab* discoveries. Three Tara Oceans MAGs classified as *Pseudo-nitzschia* (31) could be confidently placed in our phylogenomic tree (Fig. S1B). One of these contained a *dab* cluster in a MAG with high phylogenetic affinity to *P. seriata* (TARA_ARC_108_MAG_00228). This cluster is broken into two contigs, but the orientation of *dabB*/*dabC*/*dabD* strongly suggests a conserved gene structure. Additionally, a k-mer discovery tool for publicly available sequence read archives (32) suggested *dab* genes are expressed in data from various globally sampled locations. For these, we generated metaT assemblies to facilitate full-length sequence recovery. Lastly, homology searches discovered *dab* genes in the Tara Oceans Eukaryotic Gene Catalog (MATOU 1.5) and another Tara Arctic meta-assembly (33, 34).

Most of the metaT-recovered *dab* sequences exhibited high sequence identity to those insofar identified (Fig. S5). Yet, several *dab* sequences recovered from metaT data exhibited a full-length open reading frame but a separately branching position in our *dab* phylogenies. While it is not possible to directly associate these transcripts with a specific organism, we approximated their source by mapping reads from the same samples to our rDNA dataset (Fig. 3; Fig. S1A; Fig. S6). We infer that species 1, abundant at Arctic station TARA188, is *P*. cf. *obtusa* (154 raw reads from this station near-perfectly mapped to strain T5 rDNA, other reads mapped to *Fragilariopsis*), and species 2, abundant at station ARK-3M off-shore Norway (Fig. 3B) and in coastal France and Spain (Fig 3C), is *P*. cf. *plurisecta* (variable numbers of raw reads from these datasets mapped to strains Ner-D6 rDNA, Hobart5, or MLR-2013). While 12 reads from ARK-3M map to *P. delicatissima*, this species is rarely detected in other samples and shows limited genetic evidence for DA production. Whereas, *P. plurisecta* is consistently detected in samples from France and Spain, and is closely related to the *pseudodelicatissima*-like *dab* clade (Fig. S1A). The phylogenetic position of these *dab* sequences further supports the position of their inferred sources in the species tree, with *P*. cf. *obtusa* branching sister to *P. australis* and *P. seriata* and *P*. cf. *plurisecta* branching sister to *pseudodelicatissima*-like *dab* species (Fig. 2A-B; Fig. S8).

**Figure 3.**
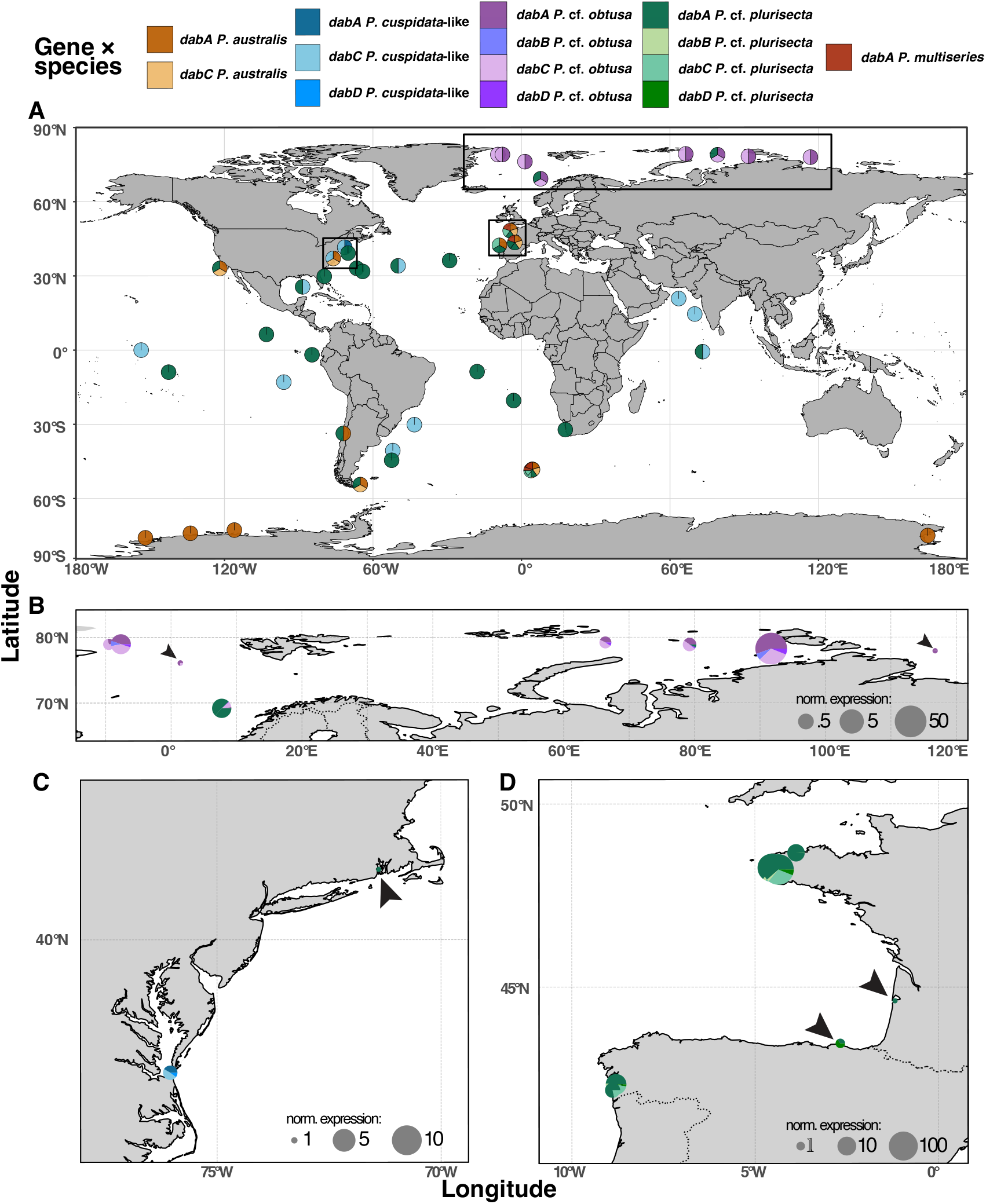
Biogeography of *dab* genes. **(A)** Boolean detection map showing the presence or absence of *dabA* and *dabC* transcripts across global metatranscriptomic datasets. Detection of *dabA* and *dabC* was observed in temperate, Arctic, and Antarctic regions. In temperate regions, *dab* gene expression was predominantly associated with *P. australis, P. cuspidata*-like, *P. multiseries*, and *P*. cf. *plurisecta*. In the Arctic, *dab* expression was associated with *P*. cf. *plurisecta* and *P*. cf. *obtusa*, whereas in the Southern Ocean, expression was associated with *P. australis*. Black rectangles mark the regions highlighted in panels B–D. **(B)** Expression of *dab* genes in the Arctic, with two dominant species inferred to be *P*. cf. *obtusa* and *P*. cf. *plurisecta*. Normalized expression is shown as reads per kilobase per million reads (RPKM). **(C)** Expression of *dab* genes along the U.S. East Coast and **(D)** in coastal regions of France and Spain. Normalized expression is shown as RPKM.

The environmental sequences exhibited low but detectable expression levels worldwide, particularly in coastal samples from France, Spain, United States Pacific and Atlantic Coast, Gulf of Mexico, the Arctic, and the Southern Ocean. In temperate regions, we predominantly observed comparable *dab* gene expression from *P. australis, P. cuspidata*-like, *P. multiseries*, and *P*. cf. *plurisecta*. In the Arctic, we observed *dab* expression from *P*. cf. *plurisecta* and *P*. cf. *obtusa*. In the Southern Ocean, we detected *dab* expression only from *P. australis*. Our analyses suggest that several unconfirmed *Pseudo-nitzschia* species may add to the complexity underlying DA production and contribute to chronic and yet under-monitored harmful algal blooms.

### Domoic acid biosynthesis dynamics in *P. pseudodelicatissima*

Given the evidence for widespread DA biosynthesis across small-celled *Pseudo-nitzschia*, we investigated the regulation of *dab* gene expression with *P. pseudodelicatissima* MT02 monoculture under silicate and nitrate limitation. Five independent replicate cultures were subjected to silicate and nitrate limitation alongside a control and repeatedly sampled over 96 hours by removing aliquots at each timepoint. Samples for transcriptomics were collected across all timepoints (n=63). Metabolomic profiling and DA quantification were conducted from 24h onward.

Silicate limitation induced growth arrest within 24h (Fig. 4A). Nitrate limitation reduced but did not halt cell division, though the difference from control was not significant (Fig. 4A). Upregulation of silicate (silicon transporter 1, *sit1*) and nitrate-starvation (high-affinity nitrate transporter 2, *NRT2*) genes validated treatment-specific transcriptional responses (Fig. S9). Transcription in this strain was driven more strongly by sampling time rather than nutrient treatment, although increasing nutrient stress over the course of the experiment likely contributed to the observed temporal patterns (Fig. S10).

**Figure 4.**
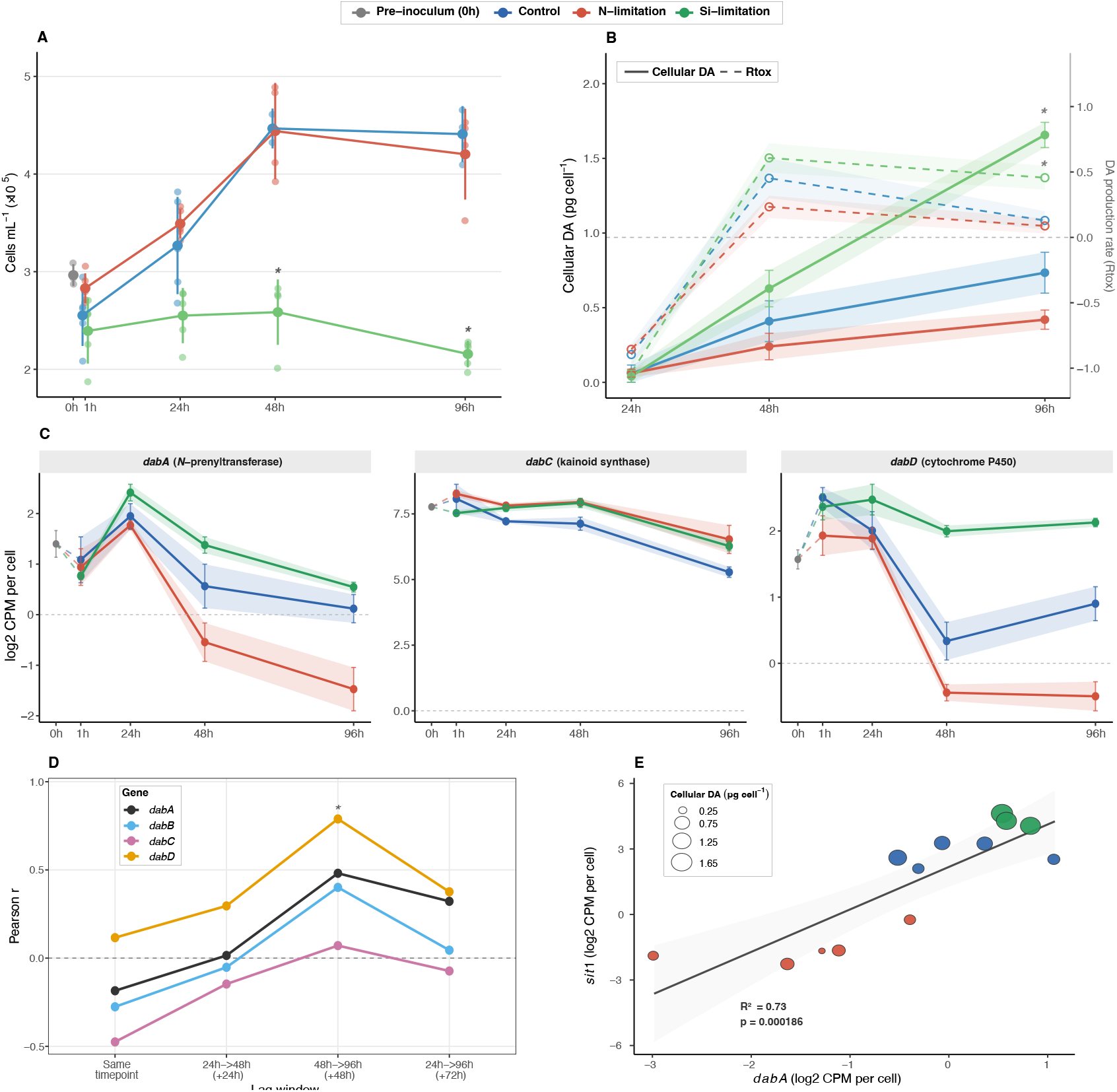
Domoic acid biosynthesis dynamics of *P. pseudodelicatissima* under nutrient limitation. **(A)** Cellular abundance (cells mL^-1^ x10^5^) over the 96 hour experiment. Lines and error bars show mean ± standard deviation (n=5 per treatment and time point). Asterisks (*) indicate pairwise Mann-Whitney U test results between Si-limitation and control treatments at the respective time points (BH-adjusted p<0.05). **(B)** Cellular domoic acid (pg cell^-1^) and domoic acid production rate (Rtox) over time. Lines and ribbons show mean ± standard error (n=2-5 per treatment and timepoint). Asterisks indicate a significant overall treatment effect on cellular DA at 96h (Kruskal-Wallis, H = 8.33, p = 0.016) and on Rtox at 96h (Kruskal-Wallis, H = 6.49, p = 0.039); pairwise comparisons did not reach significance after BH correction. **(C)** Per-cell expression trajectories of *dabA, dabC*, and *dabD* across the time course experiment. See *dabB* in Fig. S9. Lines show mean expression across biological replicates (n=5 per treatment and time point); ribbons and error bars mean ± standard error. The grey point at 0h represents pre-inoculum levels for baseline reference (n=3). **(D)** Temporal relationship between per-cell normalized transcription and toxin accumulation as shown using Pearson correlation between expression and cellular DA at different lag windows. Gene expression and cellular DA data were matched by biological replicate within treatment. Asterisk (*) indicates statistically significant predictive relationship after BH correction (BH-adjusted p < 0.05). **(E)** Co-expression of *dabA* and *sit1* expression at 96h. Points represent individual biological replicates colored by treatment; point size is proportional to cellular DA concentration (pg cell^−1^). Grey line and shaded ribbon show the linear regression fit and 95% confidence interval across all treatments pooled (R^2^=0.73, p<0.001, n=15). Si-limited samples cluster at the upper end of both axes with the highest cellular DA values, consistent with coordinated upregulation of DA biosynthesis and silicon transport under growth arrest. See all time points in Fig. S10

The *dab* genes were constitutively expressed across all conditions and time points (Fig. S11). Weighted gene correlation network analysis (WGCNA) grouped *dabA, dabB*, and *dabD* within a chloroplast-associated module (brown) enriched in methylerythritol phosphate (MEP) isoprenoid pathway genes, consistent with geranyl diphosphate being one of the substrates in the first committed step of the DA pathway. On the other hand, *dabC* was assigned to the turquoise module with broader housekeeping metabolism and regulation processes (Table S2), consistent with its two-fold higher expression relative to other *dab* genes. Differential expression analysis revealed that, under silicate limitation, *dabA, dabC*, and *dabD* expression was higher than in control cultures, with significant differences for *dabC* and *dabD* (BH-adjusted p = 0.012 and 0.001, respectively; Table S3). Under nitrate limitation, *dabA* and *dabD* were significantly repressed (BH-adjusted p = 0.001 and 0.006, respectively), while *dabC* expression was elevated (logFC = 0.89, BH-adjusted p = 0.001).

When treatments were pooled, *dabA* and *sit1* co-expression strengthened progressively over time, from R^2^=0.03 (p=0.54) at 24h to R^2^=0.54 (p=0.002) at 48h and R^2^=0.73 (p<0.001) at 96h (Fig. 4E, S11). At later timepoints, silicate-limited samples show the highest co-expression values between *dabA* and *sit1* and largest cellular DA accumulation (Fig. S12). This co-expression pattern was consistent for *dabD* and *sit1*, which shares module membership with *dabA* in the brown co-expression module. *dabB* showed no apparent relationship with *sit1* expression, consistent with its low expression variance across conditions. *dabC* displayed a negative relationship with *sit1* at all timepoints, which is in line with its assignment to the turquoise housekeeping module rather than the brown *dabABD* and *sit1*-containing module (Fig. S12).

Paired metabolomic profiling showed differences in mean cellular DA across treatments at 48h, with silicate-limited cultures accumulating 0.63 pg DA cell^−1^ compared to 0.41 pg cell^−1^ in control and 0.24 pg cell^−1^ under nitrate limitation. By 96h, mean cellular DA under silicate limitation reached 1.66 pg cell^−1^, approximately 2.3-fold higher than control and 3.9-fold higher than nitrate-limited cultures. DA production rates (Rtox, 35–37) were also highest under silicate limitation at 96h, with complete rank separation from both other treatments. Treatment comparisons performed separately within each timepoint revealed significant overall effects at 96 h for cellular DA (Kruskal-Wallis, H = 8.33, p = 0.0155) and DA production rate (H = 6.49, p = 0.0390). However, pairwise comparisons of silicate-limited cultures with other treatments did not remain significant after correction for either cellular DA or Rtox (BH-adjusted p = 0.0554; Fig. 4B). Critically, bulk total DA did not distinguish silicate-limited cultures from the control at 96h (364 vs. 347 ng mL^−1^), demonstrating that per-cell normalization is necessary to resolve the biosynthetic signal from growth-rate-driven dilution.

Primary DA biosynthesis intermediates were all detected, including *N*-geranyl-L-glutamic acid (L-NGG), 7’-hydroxy-L-NGG, 7’-carboxy-L-NGG, and isodomoic acids (Fig. S13). It should be noted that isodomoic acids A, B, and C could not be distinguished by retention time; however, based on separate analysis of this strain, isodomoic acid A was the only isomer detected (Fig. S3). Metabolomics WGCNA organized the holometabolome into discrete modules, creating a framework for integrating these results with transcriptional regulation (38). WGCNA assigned 7’-hydroxy-L-NGG, 7’-carboxy-L-NGG, and isodomoic acid A to the same coexpression module as DA. Among the sampling intervals tested, *dab* expression at 48h positively predicted cellular DA at 96h most strongly, indicating an approximately 48-hour biosynthetic lag (Fig. 4D). Only cytochrome P450 *dabD*, responsible for converting L-NGG to the 7’-hydroxy and 7’-carboxy intermediates, showed a statistically significant predictive relationship across this interval (r = 0.79, n= 13, BH-adjusted p = 0.031; Fig. 4D).

Results also show global holometabolome restructuring under nutrient limitation (Fig. S14). The silicate-limitated holometabolome was enriched for phenolic, terpenoid, and polyketide and carbohydrate metabolites (OR = 1.5–2.5, p<0.05; ; Fig. S15), including putatively annotated hydroxyphenyllactic acid, quinic acid, and p-coumaric acid-related features. On the other hand, nitrogen limitation led to enrichment of amino acid and peptide-related metabolites (OR = 4.6, p = 0.026; Fig. S15), such as putative N-acetyl-DL-glutamic acid and methionine acetaldehyde. Cross-omics integration using per-cell normalized WGCNA transcriptome and metabolome networks identified the transcriptomic brown module (containing *dabABD*) and green module (aromatic catabolism and kinase signaling) as positively correlated with the DA metabolomics module (turquoise). On the other hand, the “greenyellow” module (NRT2 nitrate transporters) and pink module (carbohydrate metabolism) were negatively correlated with the DA metabolomics module (Table S4). These pooled correlations describe coordinated transcriptional and metabolic patterns across the experiment but may also reflect shared responses to sampling time and nutrient treatment.

## Discussion

This study indicates that interpreting DA production requires differentiating biosynthetic capacity, pathway expression, and toxin accumulation, which cannot be fully resolved by species identity or discrete molecular variables. Previous research has demonstrated that DA production occurs in response to environmental conditions (2, 36, 39) and *dab* transcription can precede (11) or coincide with elevated cellular DA (10, 40–43). Here, we integrate phylogenomics, *dab* cluster characterization, environmental transcript detection, and nutrient-limited culture time-course experiments to evaluate how these lines of evidence align across *Pseudo-nitzschia*. Among the examined strains, DA production corresponded with the presence of a conserved, core biosynthesis gene cluster that is more broadly distributed across the genus than previously recognized (10, 13, 17, 18, 30), including within cryptic and understudied lineages.

The genome-resolved phylogeny of *Pseudo-nitzschia* establishes the taxonomic framework for interpreting DA biosynthesis. This approach enabled species-level resolution within the *P. delicatissima* and *P. pseudodelicatissima* complexes, clarifying longstanding unresolved relationships inferred from morphology or single molecular markers (8, 44–47). For newly sequenced strains, we directly connected the presence of the *dab* cluster to DA production in culture, showing that intact *dab* clusters corresponded with measured DA in representatives of the *P. pseudodelicatissima* and *P. cuspidata* clades. These observations highlight that small-celled taxa should not be treated as presumptively low risk or non-toxigenic. Indeed, elevated abundances of delicatissima size class strains can yield toxicities comparable to larger species (2, 9, 48). A report of low DA quotas in *P. delicatissima* (9), a species often considered baseline non-toxigenic, contrasts with the absence of *dab* genes in the representatives examined here, and further highlights the need to link toxin measurements with verified strain identity and *dab* presence. Consequently, this linked genomic and chemical context is essential for avoiding taxonomic misassignment of toxigenic potential in environmental surveys (9, 15, 49, 50).

Tracing genomic *dab* references to environmental metatranscriptomic data uncovered geographically widespread expression and a broader diversity of putative DA producers than is represented by existing genomes. While genomes identify biosynthetic potential, these environmental *dab* transcripts provide evidence of expression in natural communities. Phylogenomics, rDNA mapping, and *dab* gene phylogenies support *dab* assignments to lineages lacking genomic references, potentially representing cryptic, unsampled, or unculturable species. Moreover, these environmental *dab* transcripts implicate *Pseudo-nitzschia* taxa not routinely monitored for DA production, including *P*. cf. *obtusa, P*. cf. *plurisecta, P. pseudodelicatissima*, and a *P. cuspidata*-like group, and support existing evidence for emerging hotspots of potential DA production in the Arctic, Narragansett Bay, Gulf of Mexico, and Southern Ocean (16, 48, 51– 58). However, paired time-series measurements of species abundance, *dab* expression, and DA will be required to determine when these signals correspond to realized toxin production.

Beyond uncertainty about when *dab* expression leads to DA accumulation, regulation of the largely conserved *dab* genes across species remains poorly understood. Our genomic and biochemical analyses show that biosynthetic function can persist despite flexible locus architecture. *P. australis*, a highly toxigenic species prominent in the California Current Ecosystem (11, 59, 60), exhibits the most divergent cluster configuration among the examined lineages, with a *dabC* pseudogene co-localized with the cluster and the intact, full-length *dabC* located elsewhere in the genome. Biochemical validation of the non-clustered *P. australis* CCMP2708 DabC as an isodomoic acid A synthase demonstrates that co-localization of the *dab* genes is not required for DA pathway function. Non-clustered *dab* homologs have also recently been reported in *Chaetoceros costatus*, although their biosynthetic function remains unverified (61). Understanding how locus architecture relates to pathway regulation and downstream toxin production will require further comparative studies across *P. australis* strains and other toxigenic species.

Co-expression analysis in *P. pseudodelicatissima* begins to address this regulatory question by showing that *dab* genes do not occur in a single, coordinated transcriptional module. Association of *dabABD* with the chloroplast and MEP-isoprenoid module shows that DA biosynthesis may be coupled with precursor metabolism. Given the elevated transcript abundance of *dabC* and its coexpression with genes involved in broader metabolic functions, we hypothesize the kainoid synthase may be constitutively available, while regulation of upstream scaffold-generating steps may constrain flux through the DA pathway. The distinct transcriptional pattern and structural flexibility of *dabC*, including its relocation in *P. australis* and additional copies in *N. navis-varingica* (18, 62), motivate testing whether variation in gene architecture is associated with regulatory differences across DA-producing diatoms. Overall, this transcriptional pattern is consistent with DA biosynthesis being embedded within broader metabolic and stress-response networks (11, 40, 63, 64).

The *P. pseudodelicatissima* silicate limitation experiment connects *dab* transcription to realized chemical output, revealing how expression and cellular DA can diverge through time. Silicate limitation increased cellular DA, toxin production rates, DA biosynthetic intermediate accumulation, and *dabA-sit1* co-expression. These results indicate that elevated cellular DA reflects not only reduced growth dilution, but also increased biosynthetic activity (36, 65). Although *sit1* expression can be influenced by both Si stress and cell-cycle state (66–68), higher expression of *dabA* and *sit1* in silicate-limited samples, together with increased toxin production rates, pathway intermediates, and later cellular DA, is consistent with a response associated with Si stress. Notably, bulk DA did not consistently distinguish nutrient-limited cultures from control conditions. The relative contributions of direct *dab* regulation and upstream differences in nutrient uptake, intracellular nutrient storage, and metabolic state remain unresolved. Under the tested conditions, *dab* expression preceded cellular DA accumulation in *P. pseudodelicatissima*, aligning with field observations of delayed toxin accumulation following transcriptional activation (11, 22). In this strain, *dabD* followed by *dabA* were the strongest indicators of DA accumulation. However, genomic novelty (69) and specificity to DA producers (10, 13) make *dabA* the most informative marker of elevated DA risk. The temporal offset between *dab* expression and cellular DA accumulation further demonstrates why pathway expression should not be interpreted as equivalent to contemporaneous toxin concentration.

This study refines how genomic, transcriptomic, and chemical observations should be integrated to interpret DA risk. Relative to recent comparative studies of *dab* evolution (13) and global meta-omics surveys (15), this work contributes a genome-resolved phylogenetic framework spanning previously underrepresented diversity, biochemical validation of a functional, non-clustered kainoid synthase, and a paired expression-toxin time course under nutrient limitation. These complementary approaches highlight why no individual measurement can serve as a complete proxy for toxigenicity. Pathway presence, *dab* expression, and measured DA provide distinct evidence of genetic potential, biosynthesis activation, and realized toxin output, respectively. Recent work showing that passive sampling of copepodamide grazer cues can inform DA forecasting further highlights the value of incorporating ecological drivers alongside molecular and toxin measurements (70). Monitoring frameworks that combine these evidence types with ecological context will be better positioned to distinguish populations with DA-biosynthetic capacity from those actively producing DA and interpret the timing and magnitude of toxin risk in vulnerable ecosystems.

## Materials and Methods

Method details are available in SI Appendix. In brief, *Pseudo-nitzschia* strains were acquired from culture collections or isolated from coastal waters using standard micropipette methods (Table S1). High molecular weight DNA was extracted and sequenced using Oxford Nanopore (MinION) and/or Illumina platforms (Dataset S1). Genomes were assembled using metaFlye (71), Raven (72), or MaSuRCA (73), polished with RACON (74), and decontaminated using custom protocols. Gene prediction employed AUGUSTUS (75) or BRAKER3 (76) with functional annotation via Pfam (77), KEGG (78), and EggNOG (79) databases (Dataset S2, S3).

Phylogenomic analysis used PhyloFisher v1.1.0 (80) with 217 concatenated genes (87,978 positions) analyzed in IQ-TREE v2.3.4 (81). *Dab* gene presence/absence was confirmed by HMM (82) and BLAST (83, 84) searches and synteny analysis. Environmental *dab* sequences were identified from public metatranscriptomes using Logan-Search (32) and assembled using modified RNAseq pipelines.

*P. pseudodelicatissima* was subjected to nutrient limitation (silicate or nitrate limitation) for 96h with sampling at 0, 1, 24, 48, and 96h (n=5 biological replicates per timepoint/treatment). Weighted gene co-expression network analysis (WGCNA, 85) identified co-expression modules, with *dab* gene assignments and treatment associations assessed via module eigengenes. Culture extracts were analyzed by LC-MS/MS on an Orbitrap system. Data processing used MZmine v3.5.0 with GNPS2 molecular networking (86). Total domoic acid quantification was achieved using authentic standards (87).

## Data Availability

Supplementary Data will be available as a repository in Zenodo (10.5281/zenodo.22962408) upon publication. Raw sequence files have been submitted to NCBI GenBank under BioProjects PRJNA1532496-PRJNA1532501, PRJNA1532503, PRJNA1532507-PRJNA1532512 (Dataset 1) and will be available upon publication.

## Acknowledgments

We thank Julie A. Koester (Florida Fish and Wildlife Research Institute), Robin Sleith (Bigelow Laboratory for Ocean Sciences), and Clarissa Anderson (Southern California Coastal Ocean Observing System) for their helpful feedback and discussions. We also thank Holly Bowers (Moss Landing Marine Labs) and Vera Trainer (Olympic Natural Resources Center, University of Washington) for access to strains isolated from their laboratories. This research was supported by the National Oceanic and Atmospheric Administration (NA19NOS4780181 and NA26NOSX478C0032-T1-01 to A.E.A. and B.S.M.; NA23NOS4780293 to M.J.H, K.A.H., S.W., MERHAB publication #282), the Gordon and Betty Moore Foundation (GBMF12120 and https://doi.org/10.37807/GBMF12120 to A.M.C.-R. and P.C.D.), the National Science Foundation (OCE-2418297 to K.A.H. and A.D.) and the National Institute of Environmental Health Sciences (2P01ES028938-06 to K.H.). IJPB Institute benefits from the support of Saclay Plant Sciences-SPS (ANR-17-EUR-0007). This work was partially carried out using the URGI facilities (https://doi.org/10.15454/1.5572414581735654E12), supported by MUDIS4LS grant (ANR-21-ESRE-0048). We used computational resources of the Triton Shared Computing Cluster (TSCC) at the San Diego Supercomputer Center (SDSC; https://doi.org/10.57873/T34W2R).

