## Supplementary Information for "Genomic detection of domoic acid biosynthesis uncovers expanded toxigenic diversity across *Pseudo-nitzschia* diatoms"

**This PDF file includes:**

Supporting text
Figures S1 to S15
Tables S1 to S5
Legends for Datasets S1 to S3
SI References

**Other supporting materials for this manuscript include the following:**

Datasets S1 to S3

### Supporting Information Text

### Materials and Methods

**Strain isolation and cultivation.** Strains of *Pseudo-nitzschia* were acquired from various culture collections or isolated from coastal regions of the United States. Isolations were conducted using standard micropipette methods. See Table S1 for full details. Culture strains MT01-03, EBL58-60, NWFSC 733-4, RCC5316, CCAP1061/46 were maintained in natural seawater f/2 media (1, 2) at 16˚C under a 12:12 photoperiod. *Pseudo-nitzschia multiseries* NWFSC 713 was maintained in F/4 media at 14°C on a 12:12 L:D cycle at ~120 µmol quanta $m^{-2}s^{-1}$.Gulf of Maine *Pseudo-nitzschia* strains F, A20, and G20 were maintained in F/4 media at 14°C on a 12:12 L:D cycle at ~120 µmol quanta $m^{-2}s^{-1}$.

**DNA extraction and sequencing**. For genomes derived from *P. arctica* RCC5316, *P. arenysensis* MT, *P. australis* MT03, *P. pseudodelicatissima* MT02, *P. fraudulenta* CCAP1061/46, *P. hasleana* EBL58, and *P. pungens* var. *cingulata* NWFSC733-4, EBL59, EBL60, DNA extraction was conducted in a similar manner as described previously (3). Briefly, cultures were scaled up to 4 L of natural seawater F media and harvested via centrifugation during exponential growth (1, 2). High molecular weight DNA was extracted from up to 0.1 g of resulting cell pellets using Illustra Nucleon Phytopure Genomic DNA Extraction Kit (Cytiva). The manufacturer's instructions were followed with the following exception: the chloroform extraction and DNA precipitation steps were repeated up to 7 times in order to increase the quality of DNA. Extracted DNA was size selected using the BluePippin system with a High Pass Plus 15 kb cassette (Sage Science Cat# BPLUS03) and HMW fragment lengths verified using the Agilent 4150 TapeStation (Cat# G2992AA) with a Genomic DNA Screentape (Agilent Cat# 5067-5365).

Oxford Nanopore libraries were made with the R9.4.1 chemistry with SQK-LSK110 ligation or SQK-PSK004 PCR kit (Dataset S1) and sequenced on a MinION portable sequencing device (FLO-MIN114 flowcell). Reads were basecalled post-run with dorado v1.0.2 basecaller and the corresponding model in the “super-accurate” mode (dna_r9.4.1_e8_sup@v3.3).

For genomes derived from *P. australis* A20, *P. fraudulenta* F, *P. pungens* G20, and *P. multiseries* NWFSC 713, DNA was extracted from filtered cells using a modified CTAB method (4) as previously described (5). Extracted DNA was checked for quantity using a Qubit BR dsDNA assay and integrity on a 2% agarose gel. Illumina libraries were prepared using 1.2 µg of DNA from each isolate, sheared to 500 bp with a Covaris M220 (Covaris, Woburn, MA, USA) and cleaned with PCRclean DX beads (Aline BIosciences, Waltham, MA, USA) as input into a KAPA Hyper Prep Kit (Roche, Basel, Switzerland) following the manufacturer’s instructions. Libraries were sequenced on an Illumina NovaSeq 6000 using an S4 300 cycle kit at the University of Connecticut Center for Genomic Innovation (https://cgi.uconn.edu/).

For Oxford Nanopore libraries, DNA was enriched for fragments >1.5 kb using SPRI beads in a modified buffer (10 mM Tris-HCL, 1 mM EDTA pH 8, 1.6 M NaCl, and 11% PEG 8000). Resulting high molecular weight DNA was verified on a Fragment Analyzer before proceeding to library prep. Approximately 1 µg of high molecular weight DNA for each isolate was used for input into the Oxford Nanopore LSK114 Ligation Sequencing Kit following the manufacturer’s instructions and sequenced on a Minion using R10.4.1 flow cells. Resulting reads were basecalled with Dorado v0.4 using the super-accurate basecalling algorithm.

**Genome assembly, annotation, and phylogeny**. For genomes derived from *P. arctica* RCC5316, *P. arenysensis* MT, *P. australis* MT03, *P. pseudodelicatissima* MT02, *P. fraudulenta* CCAP1061/46, *P. hasleana* EBL58, and *P. pungens* var. *cingulata* NWFSC733-4, EBL59, EBL60, reads were adapter-trimmed by porechop v0.2.4 (https://github.com/rrwick/Porechop). For their assembly, two methods were used and compared metaFlye v2.9.3-b1797 (6) and Raven v1.8.3 (7). Assembly polishing was performed with 5 rounds of RACON v1.5.0 (8) with read mapping performed by minimap2 v2.26-r1175 (9). For each assembly, the best method was chosen based on contiguity (N50) as determined by QUAST v5.2.0 (doi: 10.1093/bioinformatics/bty266), conserved ortholog completeness as determined by BUSCO v5.7.1 (-m genome --lineage stramenopiles_odb10 --augustus_species Pseudo-nitzschia_multistriata) (8; Dataset S1).

To remove bacterial and organellar contaminants, the above datasets were subjected to the following decontamination protocol. First, organellar contigs were identified by blastn search with the following reference sequences as queries: NC_057054 *P. pungens* mitochondrial genome; NC_056171 *P. micropora* mitochondrial genome; NC_058785 *P. cuspidata* plastid genome; NC_058783 *P. pungens* plastid genome. tRNA genes were hard masked in these assemblies to avoid spurious hits. Targets with a combined coverage by query alignments higher than 50% were removed from the preliminary assemblies. Next, genes were predicted by PRODIGAL v2.6.3 (11), assuming contaminations were largely prokaryotic. The identified coding sequences were then searched in NCBI-nr using DIAMOND v2.1.9.163 blastp (--sensitive --max-target-seqs 5 --evalue 1e-5) (12). An in-house script (NRfilt, available in Supplementary Data) performed the taxonomic polling assessment based on the tab-separated output of DIAMOND, with decontaminated contigs and gene models as output. Briefly, the process assigns taxonomy to the best hit per query using NCBI resources, and aggregates hits at the scaffold level to decide whether each scaffold is an ingroup (Bacillariophyta) or contaminant. High-confidence hits are defined by thresholds on query coverage (>50%) and percent identity (>75%), and scaffolds are classified primarily by whether a sufficient fraction of their genes' high-confidence hits belong to predefined ingroup or to contaminant groups. If no decisive high-confidence signal exists, the script falls back to low-confidence evidence and applies more permissive rules, ultimately retaining scaffolds unless contaminant signals clearly dominate. Finally, gene models in genomes were predicted with augustus v3.5.0 (13) or BRAKER3 (14) (Dataset S1 for details).

For *P. australis* A20, *P. fraudulenta* F, *P. pungens* G20, and *P. multiseries* NWFSC 713, several hybrid [MaSuRCA v4.1.0 (15), Wengan v0.2 (16), and SPAdes v3.15.5 (17) with LongStitch v1.0.5 (18)] and long-read assemblers [Shasta v0.11.1 (19) and Flye v2.9.2 (6)] were tested and evaluated with bbmap v39.05 and completeness with BUSCO v5.5.0, choosing the assemblies based on BUSCO completeness and contiguity. For most genomes, MaSuRCA provided the highest quality assemblies, except for *P. australis* A20 for which Wengan performed better (Dataset S1). Bacterial contamination was removed using NCBI's Foreign Contamination Screen tool, while mitochondrial and chloroplasts were removed using blastn searches against a curated organelle reference, including mitochondrial (NC_027265.1 *P. multiseries*, NC057054.1 *P. pungens*, NC056171.1 *P. micropora*, NC_056161.1 *P. cuspidata*, NC060304.1 *P. delicatissima*) and chloroplast (NC_027721.1 *P. multiseries*, PQ010277.1 *P. sabit*, PQ010276.1 *P. galaxiae*, PQ010275.1 *P. brasiliana*, NC058783.1 *P. pungens*, NC058785.1 *P. cuspidata*) references retrieved from NCBI. For gene model prediction, RNAseq data from *P. delicatissima*, *P. australis*, *P. arenysensis*, and *P. granii* (PRJNA248394, PRJNA432444, PRJNA903882) were retrieved as evidence and mapped to the reference genomes using HISAT2 (20). Mapping results from all samples were then combined into a single BAM file. Repeat annotation and masking was peformed with RepeatModeler (v1.89.2; https://www.repeatmasker.org/RepeatModeler/) and gene model prediction was performed on the soft-masked reference genome with mapped RNAseq data as evidence using BRAKER3 v3.0.8 (14) with Augustus v3.4.0 (13).

For pre-existing transcriptomic data (Dataset S2), TransDecoder.LongOrfs v5.7.0 (21) was used for coding sequence extraction.

To provide a comparable input for our Pfam enrichment analyses, versions of *Nitzschia navis-varingica* genomes were decontaminated as above and their proteomes predicted with augustus v3.5.0 (13).

Functional annotation followed employing common protein domain databases Pfam v37.0, PANTHER v18.0, TIGRFAM v15.0, KEGG v29-10-2024, and EggNOG v5 using a combination of tools (diamond v2.1.9, eggNOG-mapper v2.1.10, Interproscan v5.69-101.0, kofamscan v1.3.0 (12, 22–25).

Phylogenomic analysis was performed using PhyloFisher v1.1.0 (26) and included newly sequenced as well as pre-existing datasets (Dataset S2; (3, 27–32). A concatenated alignment of 217 genes (87,978 positions, 28,641 parsimony-informative, 13,996 singleton, and 45,340 constant sites) was used as input for IQ-TREE v2.3.4 (33) to first generate a guide tree under the ELM+C60+G model, and a final tree under the Posterior Mean Site Frequency model and with a maximum of 1000 ultra-fast bootstrap replicates (34, 35). Alignments and newick-format trees provided in Supplementary Data.

Strain identification was performed using a previously described ITS2 dataset (36), except here the entire region spanning the V9r binding site to the 28SR binding site was utilized (includes fragments of 18S and 28S on both ends, and full-length ITS1, 5.8S, and ITS2, roughly between 1510R and 28SR primer binding sites (37, 38)). Note that only sequence fragments were found for *Pseudo-nitzschia australis* CCMP2708 and *Pseudo-nitzschia multiseries* CLN-47, whereas no sequences were found in MAGs and *Pseudo-nitzschia multistriata* B856. Sequences were aligned with Muscle5 v5.1 (39), manually edited to resolve misaligned positions; the final alignment had 1,678 positions, 667 parsimony-informative, 105 singleton, 906 constant. Tree was inferred with IQ-TREE v2.3.4 under the best-scoring model based on Bayesian Information Criterion (GTR+F+R4). Alignments and newick-format trees provided in Supplementary Data.

*Dab* cluster gene structures were found and manually curated in genome assemblies using the BLASTN and annotation functions of Geneious Prime v2025.2. Open reading frames were extracted, aligned by gene using MUSCLE 5.1, and used for CDS phylogenies using IQ-TREE v2.4.0.

Diatom images were three-dimensionally modeled in Blender 4.2 (Blender Foundation, Amsterdam, Netherlands) based on morphological illustrations from Hasle and Syvertsen (1997) (40). Cell geometries were constructed using box modeling techniques, with frustule ridge patterns created using hand-drawn height maps for surface displacement. Texturing was done procedurally using Blender's shader node system. Final images were rendered using the Cycles (part of Blender) path-tracing render engine with physically-based lighting.

**Gene enrichment analysis**. Rapid protein family evolution was assessed by CAFE5 v5.1.0 (41). Pfam or PANTHER domain counts were extracted from genome assembly annotation tables (transcriptome datasets were excluded) and merged, then Pfams with <50% occupancy across the entire dataset were removed; these concerned 1544 singletons and Pfams with no obvious phylogenetic pattern, i.e., not shared between species of the same lineage. For the CAFE5 analysis, large families were separated by clade_and_size_filter.py (default parameters). The final phylogenomic tree was pruned to remove branches that were excluded, and then converted to an ultrametric tree using r8s v1.80 (42), placing the common ancestor of *Phaeodactylum tricornutum* and *Pseudo-nitzschia multistriata* to 108 Mya (i.e., median divergence time according to 40). Three CAFE5 runs were performed with: 2 gamma categories plus Poisson distribution for the root frequency distribution, and checked for convergence. Consensus results, i.e. where direction of a gene family's accelerated evolution at a given node was identical for the three runs, were plotted with a custom code available as a jupyter-notebook (Supplementary Data).

**Global metatranscriptomic data search and assembly.** Public metatranscriptomic datasets were queried for *dab* genes using the Logan-Search tool, which is a k-mer-based search engine of the Sequence Read Archive (44). Datasets were then selected based on strength of the hit and geographic location and accessed via NCBI SRA (45) .

MetaT assemblies were generated using a modified RNAseq Annotation Pipeline (46). Briefly, raw reads were adapter- and quality-trimmed using fastp v0.23.2 (trimmomatic option; 44, 45). rRNA-matched trimmed were further filtered out using BBduk (https://jgi.doe.gov/data-and-tools/software-tools/bbtools/; rDNA databases PR2 v4.12.0, RFAM v14.1, SILVA v138). Transcriptomes were assembled using MEGAHIT v1.2.9 (49), then combined at 95% sequence identity using linclust of mmseqs2 release 14 (--min-seq-id 0.95 -c 0.8 --cov-mode 1 --cluster-mode 2) (50). Open reading frames were determined and translated by GeneMarkS v5.1 (doi: 10.1093/nar/29.12.2607). The Arctic metatranscriptomic dataset was adopted from Füssy et al. (2025) (51). Read mapping rates were obtained by mapping raw reads to contigs or open reading frames by hisat2 v2.1.0 (20) (Sequence Read Archives used: TARA188: ERR6349669, ERR6349672, ERR6349680; Norway offshore: SRR8393572).

To identify environmental homologs of the *dab* genes, profile hidden Markov models (HMMs) were used built with reference protein sequences compiled from *Pseudo-nitzschia* and *N. navis-varingica* genomes and transcriptomes. These were supplemented with characterized homologs found in red algae. Reference sequences were aligned using MAFFT v7.256 (52) and HMMs constructed using HMMER v3.32 (53).

The metatranscriptomic protein assembly was used for HMM queries. Candidate hits were retained if they met domain-level criteria (evalue<1e-10 and 0.6 coverage) to ensure sequences contained motifs characteristic of *dabA* and *dabC*. To assign HMM-derived candidates to specific samples, regions, and assemblies, these hits were subsequently used as query sequences in combination with reference genes in MMseqs2 (50) to consolidate homologs and minimize redundancy across environmental datasets. MMseqs2 results were filtered to retain the best-scoring hit per query based on e-value (<1e-5), coverage (>0.5), and results from additional HMMsearch.

The resulting high-confidence environmental *dabA* and *dabC* sequences were incorporated into phylogenies with reference sequences to confirm identity and species-level designation. Protein sequences were aligned using MAFFT v7.256 and analyzed using maximum-likelihood phylogenies generated with IQ-TREE v2.4.0. Best-fit substitution models were selected independently for each alignment using ModelFinder according to BIC, namely JTT+I+G4 for *dabA* and VT+G4 for *dabC*. Branch support was assessed using 1,000 ultrafast bootstrap and SH-aLRT replicates (33). Outgroups were selected based on previous biochemical characterization (54–56). Sequences that clustered with *dabA* and *dabC* clades were considered true environmental homologs. Other sequences with long-forming branches or ambiguous placements were not included in downstream visualization.

**Metabolite extraction from *Pseudo-nitzschia* species.** For strains, *P. arenysensis* MT01, *P. pseudodelicatissima*, *P. australis* MT03, *P. hasleana* EBL58, *P. pungens* ELB59 and EBL60, *P. arctica* RCC5316, *P. fraudulenta* CCAP1061/46, and *P. pungens* NWFSC 733 and 734, metabolites were extracted using solid-phase extraction (SPE) with Agilent Bond Elut PPL cartridges (200 mg sorbent, 3 mL volume), following an approach adapted from established dissolved organic matter workflows (57, 58). During exponential growth, 50 mL aliquots of culture were collected and acidified to pH 2 using concentrated HCl. Prior to sample loading, cartridges were conditioned sequentially with LC–MS grade methanol and acidified LC-MS grade water (pH 2). Acidified culture samples were applied to the cartridges under vacuum at a slow, dropwise flow rate to retain metabolites from both dissolved and particulate phases. Following loading, cartridges were rinsed with acidified LC–MS grade water (pH 2) to remove residual salts. Sorbent beds were dried under a stream of nitrogen gas, and retained compounds were eluted with 2 mL LC-MS grade methanol. Eluates were concentrated to dryness using a SpeedVac and reconstituted in 100 µL of 80% (v/v) aqueous methanol containing 0.1% (v/v) formic acid. Reconstituted extracts were stored at −80 °C prior to LC-MS analysis.

For strains *P. multiseries* NWFSC 713, *P. australis* A20, *P. pungens* G20, and *P. delicatissima* D, cultures containing between 479,600 - 3,105,000 cells were filtered ono HVLP (0.45 µm) or PCTE (0.2 µm) filters. Filters were stored at -80˚C until extraction. Extraction was performed using 50% aqueous methanol (LC-MS grade), sonicated for 10 minutes, and vortexed for 5 minutes. Filters were then removed and resulting extracts were pelleted and spin-filtered (0.22 µm) to remove debris. Samples were dried under vacuum and resuspended in 400 µL of 80% (v/v) aqueous methanol containing 0.1% (v/v) formic acid. Samples were stored at −80 °C prior to LC-MS analysis.

**Biochemical characterization of *P. australis* kainoid synthase enzyme**. *P. australis* CCMP2708 kainoid synthase DabC was heterologously expressed in *Escherichia coli* and purified as previously described (3, 59, 60). The gene was codon-optimized for *E. coli* expression, domesticated for SapI and BsaI cut sites, and designed with an N-terminal His-6 affinity tag in pET-28a(+) vector and ordered from Twist Bioscience.

Plasmid constructs were introduced into chemically competent *E. coli* BL21(DE3) cells and selected on LB agar containing kanamycin (50 µg $mL^{-1}$). Individual transformants were used to initiate overnight starter cultures, which were subsequently diluted into 500 mL Terrific Broth supplemented with 4% (v/v) glycerol and grown at 37 °C with shaking to mid-log phase (OD-600 ≈ 0.6). Cultures were cooled on ice prior to induction with 1 mM isopropyl β-D-1-

thiogalactopyranoside (IPTG), then incubated at 18 °C with shaking for approximately 18 h. Cells were collected by centrifugation, flash frozen, and stored at -80 °C until purification.

Cell pellets were thawed on ice and resuspended at 5 mL buffer per 5 mg wet weight in lysis buffer (10 mM HEPES, 100 mM NaCl, 25 mM imidazole, 0.2 mM DTT, 2.5 mM EDTA, 20% glycerol, pH 7.5) supplemented with lysozyme (1 mg $mL^{-1}$). Cells were disrupted by sonication (Qsonica probe; 50% amplitude, 15 s on/45 s off cycles, 7 min total processing time). DNase I (5 µg $mL^{-1}$ final concentration) and $MgCl_2$ (5 mM) were added midway through sonication to reduce viscosity. Insoluble material was removed by centrifugation at 40,000 × g for 30 min at 4 °C, and the clarified lysate was passed through a 0.45 um syringe filter prior to downstream purification.

Purification was performed using immobilized metal-affinity chromatography purification of His6-tagged protein using a 5 mL HisTrap FF column (Cytiva). The column was pre-equilibrated with lysis buffer and cleared lysated was loaded onto it via peristaltic pump. The column was then washed with 50 mL wash buffer 1 (10mM HEPES, 100 mM NaCl, 25 mM imidazole, 10% glycerol, pH 7.5), followed by 50 mL of wash buffer 2 (10mM HEPES, 100 mM NaCl, 50 mM imidazole, 10% glycerol, pH 7.5), while collecting fractions of the flowthrough. The target protein was eluted using 5 mL of elution buffer (10mM HEPES, 100 mM NaCl, 500 mM imidazole, 10% glycerol, pH 7.5). Fractions were analyzed by SDS-page. Fractions containing the target protein DabC were pooled and buffer-exchanged using a PD-10 column (Sephadex G-25 M, Cytiva) that was pre-equilibrated with storage buffer (50 mM HEPES, 250 mM NaCl, pH 8, 10% glycerol). DabC protein fractions were concentrated with Amicon Ultra-15 centrifugal filters, aliquoted, flash frozen in liquid nitrogen, and stored at -80˚C until further analysis. The protein was quantified using denatured protein UV absorbance at 280 nm at multiple dilutions with Milli-Q water.

The DabC enzyme activity assay was also performed as previously described (3). A reaction mixture was prepared in a final volume of 100 µL containing 100 mM HEPES (pH 8.0), 100 mM KCl, 10% glycerol, 1 mM L-ascorbate, 6.25 mM 2-oxoglutaric acid, 25 µM to 1 mM of either 7′-COOH-L-NGG, 50 µM DabC, and 50 µM $FeSO_4 \cdot 7H_2O$, and incubated at 25°C overnight (~15 h). The reaction was quenched by adding 100 µL of methanol, followed by centrifugation. The supernatant was purified via filtration with a nylon 0.22 µm pore CA membrane (Costar Spin-X).

DA standard was purchased from the National Research Council of Canada (61). Preparation of all non-commercial substrates was used as prepared in previous studies (59, 62). All substrates and enzymatic assay products were verified using retention time and mass via liquid chromatography–mass spectrometry.

Liquid chromatography-mass spectrometry (LC-MS) analyses were performed following an established workflow adapted from prior work (3, 60). Samples were analyzed on an Agilent Technologies 1200 Series system with diode array detector coupled to an Agilent Technologies 6530 accurate-mass QTOF LC-MS. Separation was achieved using reversed-phase chromatography on a Phenomenex Kinetex C18 column (5 µm particle size, 100 Å pore size, 150 × 4.6 mm). The mobile phases consisted of water containing 0.1% (v/v) formic acid (solvent A) and acetonitrile containing 0.1% (v/v) formic acid (solvent B). Elution was carried out at a flow rate of 0.75 mL $min^{-1}$ using the following gradient program: an initial hold at 5% B for 1 min, a linear increase to 35% B over 30 min, a ramp to 100% B over 1 min, a 1.5 min hold at 100% B, re-equilibration to 5% B over 2.5 min, and a final hold at 5% B for 2 min.

***P. pseudodelicatissima*** **gene expression and metabolomic patterns during nutrient limitation**. To characterize domoic acid biosynthesis gene expression patterns of *P. pseudodelicatissima* strain MT02 under nutrient limitation, *P. pseudodelicatissima* cells were pelleted during exponential growth phase and resuspended in three conditions: control (F/2+Si media), F/2 without silicate (“SiLim”), F/2+Si without nitrate (“NiLim”), and grown for 96 hours. Cultures were sampled at five timepoints over 96 hours: immediately prior to resuspension (Ti, 0h; pre-experiment inoculum), 1 hour after resuspension (T0, 1h; treatment establishment), and at 24h (T1), 48h (T2), and 96h (T4) post-resuspension. Each treatment and time point was conducted with five biological replicates. Cellular abundance and metatranscriptomics samples were collected at each time point and condition (n=63). Metabolomic data was collected beginning at 24h (n=39). However, sampling coverage was partial at 24h (Control: 5/5; NiLim:

3/5; SiLim: 2/5) and at 96h (Control: 5/5; NiLim: 5/5; SiLim: 3/5) due to sample loss. A single pre-experiment inoculum sample (Ti, 0h) was included for metabolomic analysis as a baseline reference.

Cellular abundance of each biological replicate was determined using an inverted light microscope (Olympus CKX53) with a Neubauer improved hemocytometer (Marienfeld, CAT No. 0650030). Each sample was counted in duplicate and cellular abundance was calculated using the average cell count total.

Samples for whole culture metabolomics were processed in a manner identical to previously described (see "Metabolite Extraction from *Pseudo-nitzschia* species"). In this case, 30 mL of whole culture was processed using solid-phase extraction (SPE) with Agilent Bond Elut PPL cartridges (200 mg sorbent, 3 mL volume), following an approach adapted from established dissolved organic matter workflows (57, 58).

DA production rate was calculated as Rtox, which is the change in total DA concentration (cellular abundance multiplied by total DA content per cell) between two time points, divided by the logarithmic mean cell concentration over that interval (63–65). Treatment differences in cell density, cellular DA, and DA production rate (Rtox) were assessed separately at 24, 48, and 96 h using Kruskal-Wallis tests. Pairwise treatment comparisons used Mann-Whitney U tests, with Benjamini-Hochberg correction across the three comparisons within each timepoint and metric. Temporal differences within treatments were assessed using Friedman tests on complete replicate time courses and paired Wilcoxon signed-rank tests matched by replicate, with Benjamini-Hochberg correction across timepoint comparisons within each treatment and metric.

***P. pseudodelicatissima* transcriptomics data processing.** Transcript abundance estimates (transcripts per million, TPM) were obtained for all predicted open reading frames downstream analysis to adjust for sequencing depth and transcript length. Expression values were $\log_2$ transformed prior to analysis. To reduce noise and computational complexity, genes were ranked by variance across samples, and the 2,000 most variable genes were retained for network construction. Transcripts encoding *dab* genes (*dabABCD*) were explicitly retained regardless of variance to ensure their inclusion in the analysis.

Weighted gene co-expression network analysis (WGCNA) was performed using the WGCNA R package (66) clustering based on Euclidean distance was used to assess potential outliers. A co-expression network was constructed using a signed hybrid topology and a soft-thresholding power selected to approximate scale-free topology (power = 12). Modules were identified using dynamic tree cutting with a minimum module size of 20 genes. Module membership for individual genes was quantified using eigengene-based connectivity (kME). Domoic acid biosynthesis genes (*dabABCD*) were assigned to modules and evaluated for hub-like behavior based on kME values. Associations between module eigengenes and experimental variables were assessed using Pearson correlation, with significance evaluated using the Benjamini-Hochberg p-values. Functional annotations included KEGG, KOfam, PFam, KOG, and UniProt descriptors, and functional enrichment patterns were summarized descriptively based on the frequency of annotation categories within modules.

To evaluate whether bulk TPM normalization confounded treatment effect interpretation, a parallel per-cell normalization analysis was conducted as the primary analysis. Raw transcript counts were divided by cell density (cells $mL^{-1}$) at each sampling timepoint, scaled to per $10^5$ cells, and TMM-normalized using edgeR (67) to correct for residual library size differences. The resulting matrix was log2-transformed and used for all primary WGCNA and differential expression analyses. Due to strong experimental structure in the data, scale-free topology could not be achieved at the conventional $R^2 \geq 0.85$ threshold; soft-thresholding power was therefore selected empirically (power = 6, signed hybrid network) based on minimizing unassigned genes and maximizing consistency of *dab* gene module assignments with the TPM network. *dab* gene module assignments were identical between normalization approaches (dabA, dabB, dabD: brown module; dabC: turquoise module), while treatment-eigengene correlations differed substantially for several modules ($|\Delta R| > 0.5$ for green and magenta modules), consistent with growth-rate confounding in bulk TPM data. Cases where normalization methods affected biological interpretation are noted explicitly in the results.

Differential expression was assessed using limma (68) with empirical Bayes moderation. An additive linear model was fitted across T1–T4 to estimate mean treatment effects versus control after accounting for timepoint. A separate interaction model was used to identify timepoint-specific treatment effects, with within-timepoint contrasts extracted for biological interpretation. All reported p-values are Benjamini-Hochberg adjusted unless otherwise noted. DESeq2 (69) was used complementarily with a factorial design and likelihood ratio test to identify genes with significant time-dependent treatment effects. Differentially expressed gene sets were summarized functionally using KEGG, KOfam, and PFam annotations.

***P. pseudodelicatissima* metabolomic data acquisition and processing.** Untargeted LC-MS/MS acquisition was performed on a Vanquish Flex UHPLC system coupled with an Orbitrap Exploris 240 (Thermo Fisher Scientific, Bremen, Germany) operating in positive mode. Chromatographic separation was performed on a Kinetex C18 reversed phase column (1.7 µm, 100 Å, 150 x 2.1 mm; Phenomenex, Torrance, CA) using a 15 minute gradient at a flow rate of 0.5 $mL^{-1}$. Mobile phase A consisted of water with 0.1% formic acid, and mobile phase B consisted of acetonitrile with 0.1% formic acid. All solvents were LCMS grade. The gradient was programmed starting at 5% B at 0.5 min, increased linearly to 50% B at 8 min, ramped to 99% B at 10 min and held until 12 min, followed by re-equilibration to 5% B by 13 min and held until 15 minutes. Blanks were run every 10 samples to ensure consistency.

Full MS scans were acquired in the Orbitrap at 240,000 resolution over a range of m/z 100–1000. Data-dependent acquisition (Top10) was used to select the most intense precursor ions per cycle for fragmentation. MS/MS spectra were acquired at 15,000 resolution with a 1.0 m/z isolation window and stepped normalized higher-energy collisional dissociation (HCD) energies of 20-50%. See full details using the MassIVE dataset MSV000099945 (70).

To quantify total domoic acid (tDA) in culture samples, a standard curve was generated through concurrent analysis, spanning six orders of magnitude (10,000 - 0.1 ng $mL^{-1}$) of DA dilutions to establish an integrated peak area range. Quantitative analysis was then conducted using the GNPS2 Dashboard (71) using the extracted ion chromatogram m/z 312.144 (± 5 ppm, retention time 3.6-3.9 min) for the $[M+H]^+$ DA ion. Detection of DA, isodomoic acids A-C, N-geranyl-L-glutamic acid, 7'-carboxy-N-geranyl-L-glutamic acid, and 7'-hydroxy-N-geranyl-L-glutamic acid features were also achieved using feature-based molecular networking (71, 72) and subsequently confirmed by aligning the retention time with that of concurrently run synthetic standards. As previously noted, the DA standard was obtained from the National Research Council of Canada (61), and non-commercial intermediates were prepared as previously described (59, 60, 62).

Raw data files were converted to mzML for downstream processing using MZmine v3.5.0 (73). Mass detection was performed using noise thresholds of $1.0\times10^4$ for MS1 scans and $1.0\times10^3$ for MS2 scans, with a minimum feature height of $1.0\times10^3$. Chromatogram building used a scan-to-scan m/z tolerance of 10 ppm with a minimum of 5 consecutive scans. Chromatographic smoothing and stable ionization correction across samples were enabled. Feature resolution used an approximate peak full width at half maximum of 8 seconds with an intra-sample m/z tolerance of 0.0015 m/z or 5 ppm and an intra-sample retention time tolerance of 0.08 minutes. Feature alignment was performed using a sample-to-sample m/z tolerance of 10 ppm and a retention time tolerance of 0.40 minutes. Gap filling was performed using default parameters, and features detected in fewer than 3 samples were removed from the aligned feature list. The resulting feature table and corresponding MS2 spectra were exported for FBMN using GNPS2 (72, 74). Spectral library matching against the GNPS LC library was performed with a minimum cosine score of 0.70, a minimum of 6 matched fragment peaks, a fragment ion tolerance of 0.05 Da, and a precursor mass tolerance of 0.02 Da, retaining the top 1 library match per feature. See details of FBMN job here:
https://gnps2.org/status?task=a18ce557643a4b57b157bf0bb4988164#.

The FBMN feature quantification table was further analyzed in RStudio v2024.12.1 using WGCNA, differential abundance, and enrichment analysis (75). Samples corresponding to media blanks, extraction blanks, and LCMS run blanks were excluded from downstream analysis after applying blank exclusion during data normalization in R. Features with missing values in more than 50% of samples were removed, and remaining missing values were imputed with half the feature minimum. Abundances were log-transformed and normalized using probabilistic quotient normalization (PQN) to correct for variation in extraction efficiency. A bottom 10% variance filter was applied. A cell-count-normalized matrix was also constructed by dividing raw feature abundances by the corresponding cellular abundance measurement prior to log transformation.

To distinguish genuine treatment-driven metabolome changes from those attributable to differential growth rates between treatments, a parallel cell-count-normalized matrix was constructed. Raw feature abundances were divided by the corresponding cell density measurement (cells $mL^{-1}$) at each sampling timepoint prior to log2-transformation, producing per-cell abundance estimates. This normalization approach addresses the growth-rate confound inherent in bulk metabolomics normalization, where differences in cell density between treatments can obscure or create apparent abundance differences that do not reflect genuine per-cell metabolic regulation. Linear mixed-effects models (LMM) with replicate as a random effect were used to test for treatment main effects and time × treatment interactions on module eigengenes under both PQN and per-cell normalization, using likelihood ratio tests with Benjamini-Hochberg adjustment. Treatment effects that were significant only under per-cell normalization were interpreted as reflecting genuine per-cell metabolic regulation masked by growth-rate differences in the PQN data; treatment effects significant in both normalizations were considered robust.

A WGCNA co-expression network was constructed using a soft-thresholding power selected to approximate scale-free topology (power = 6; R2 > 0.85) (66). Modules were identified using the blockwiseModules function with a signed hybrid network type, signed Nowick TOM, minimum module size of 30 features, and a merge cut height of 0.35 to combine highly similar modules. Module eigengenes from the metabolomics WGCNA and the metatranscriptomics WGCNA were aligned to shared samples. Pearson correlations were computed between all pairwise combinations of metatranscriptomics and metabolomics module eigengenes using pairwise complete observations. P-values were computed using Student's t-test and adjusted for multiple comparisons using the Benjamini-Hochberg method. Cross-omics correlations pooled samples across timepoints and treatments without adjustment for time, treatment, or repeated sampling of individual cultures and were interpreted as exploratory associations.

Pairwise treatment contrasts were performed at each timepoint using Welch's t-tests on cell-count-normalized feature abundances. Features were considered differentially abundant at nominal $p < 0.05$ and $|log2FC| > 1$. Chemical pathway annotations from the NPClassifier framework were collapsed into seven broad categories. Enrichment of annotated chemical classes among differentially abundant features was assessed using one-sided Fisher's exact tests relative to the background frequency in the full feature matrix, with results pooled across timepoints per comparison type and adjusted using the Benjamini-Hochberg method.

### Figures

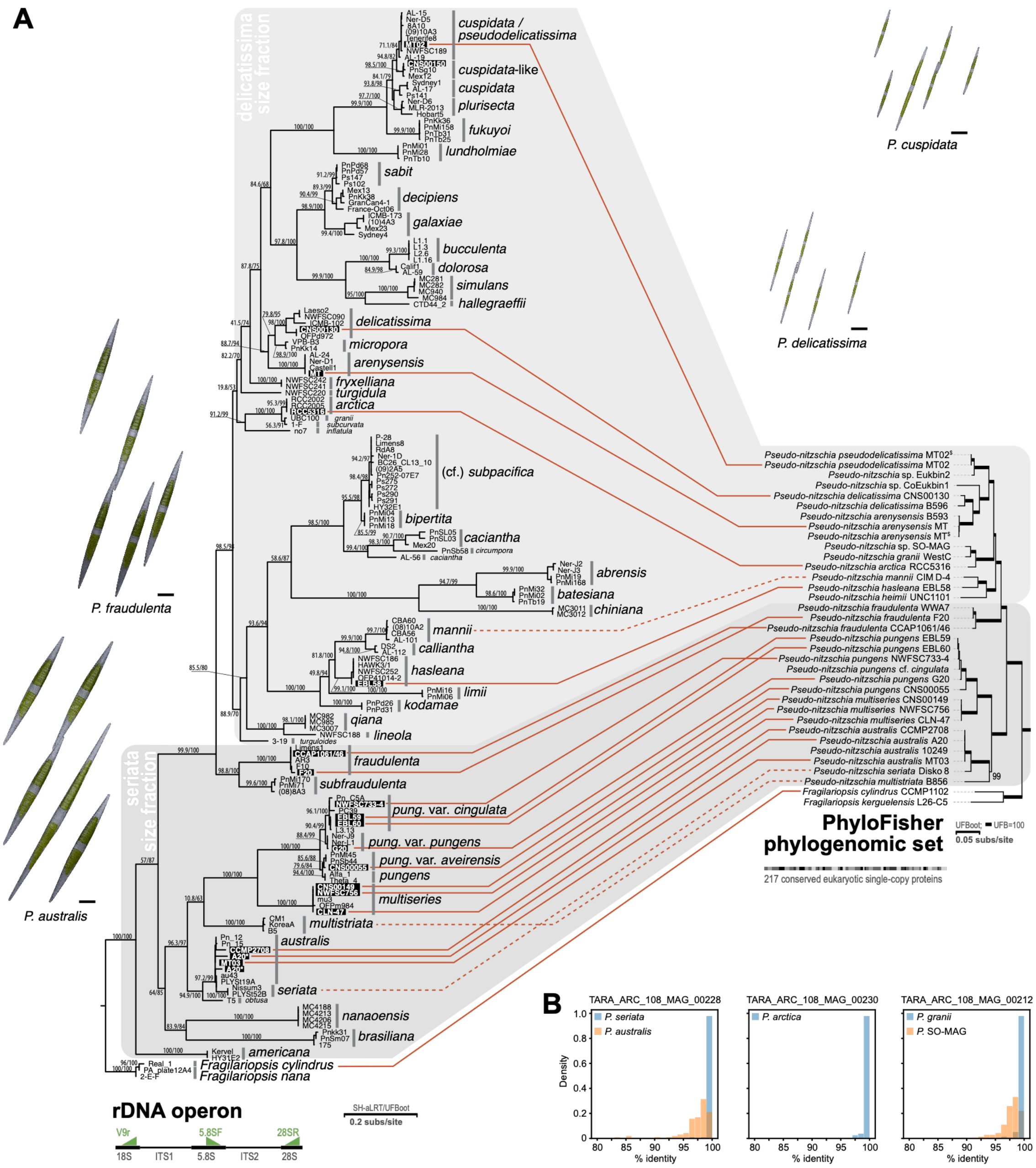

**Fig. S1.** The *Pseudo-nitzschia* genomic resource phylogeny. (A) Side-by-side comparison of ribosomal DNA operon (left) and phylogenomic (right) phylogenies. Full lines link data from the same strain, whereas dashed lines link data from the same species. Asterisks mark two different rDNA sequences extracted from *P. australis* strain A20. In previous works, ITS1/2 data offered better diagnostic resolution for taxa, but lacked backbone support. Here, we compare the ribosomal operon ML phylogeny (1,678 positions spanning the 1520R and 28SR primer binding sites and the ITS1/5.8S/ITS2 regions between, GTR+F+R4, branch support expressed as ultra-fast bootstrap/SH-aLRT values of 1000 replicates each) with that of the phylogenomic dataset (217 proteins, 87,978 positions, ELM+C60+G guide tree, PMSF final tree, branch support expressed as ultra-fast bootstrap values for a maximum of 1000 replicates, see Methods). Note the resulting topologies are highly congruent and crucial branchings highly supported. The phylogenomic tree is identical to that in Fig.1, except that two nodes were rotated to better illustrate this congruence (node rotation does not affect branching order). Diatom images decorating this panel illustrate inter-species size differences and intra-species size variability, with scale bars representing 10 µm. The images were box-modeled in Blender based on illustrations from Hasle and Syvertsen (1997), textured procedurally, and rendered in Cycles (see Methods). (B) Pairwise sequence identity of the above phylogenomic markers found in three *Pseudo-nitzschia* MAGs. In our pilot single-gene phylogenetic assessment, we noticed various levels of chimeric origin for these MAGs. For the three MAGs shown, phylogenetic noise was smaller, and they were consistently branching with one or two species from the reference set. Pairwise identity with these closest relatives is shown as histogram densities. The inferred origin of these MAGs was the one with the 100% identity peak.

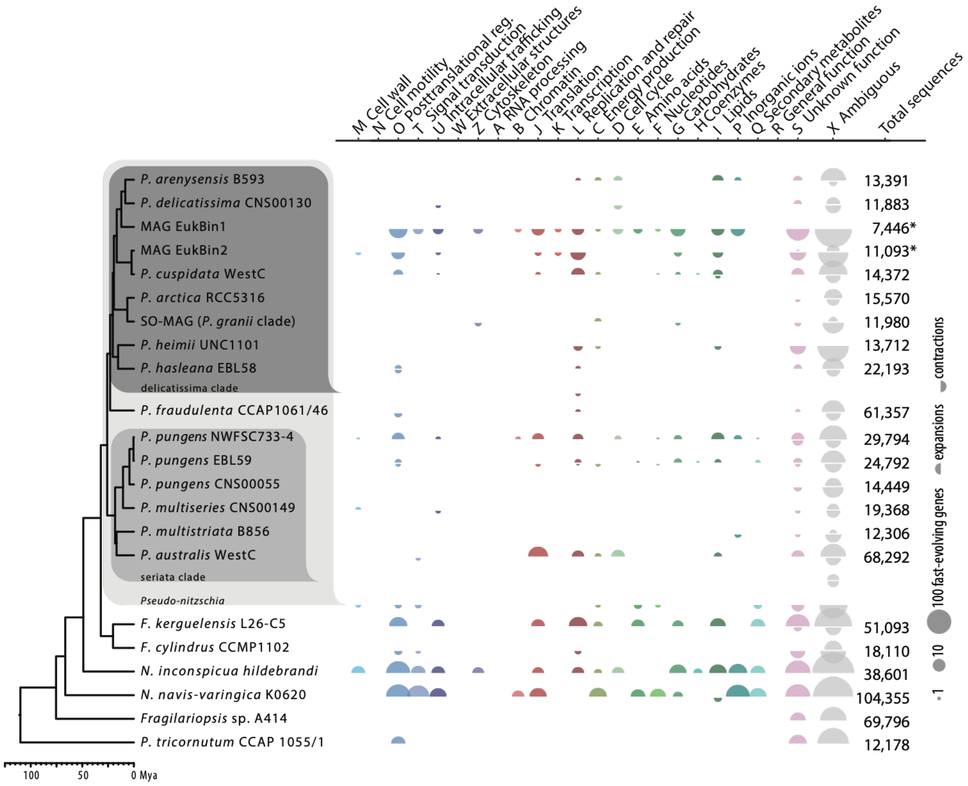


**Fig. S2.** Accelerated evolution of protein families in *Pseudo-nitzschia*. Summary of the number of genes found in protein families with rapidly expanding or contracting evolution, calibrated by phylogenetic distance (CAFE5). The analysis found little evidence for accelerated evolution at the ancestral node of major *Pseudo-nitzschia* clades, suggesting species radiation is not associated with significant functional expansion. Large families excluded from analysis. * – rapid contraction likely an artifact of incomplete genomes (MAGs).

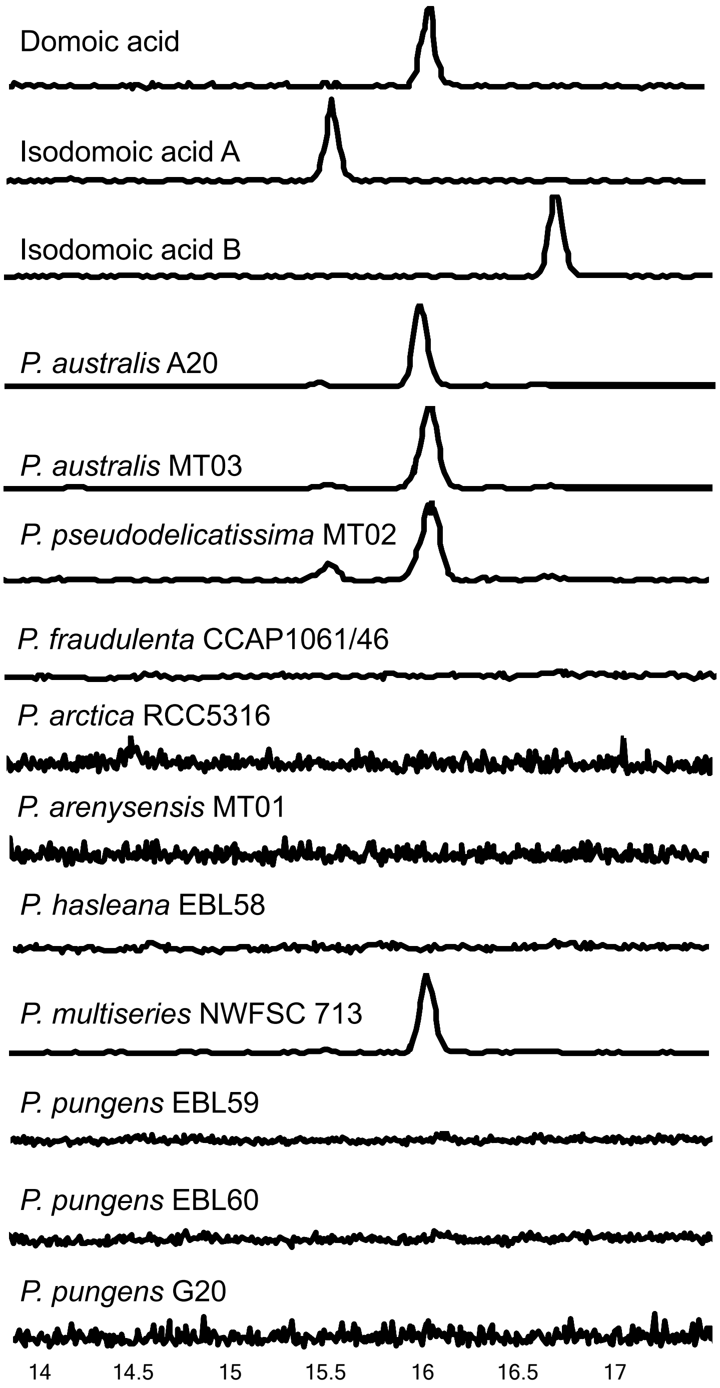


**Fig. S3.** Extracted ion chromatograms for domoic acid (310.1 m/z) and isodomoic acids A and B in comparison with standards. Isodomoic acids B and C have been shown to co-elute using our methods.

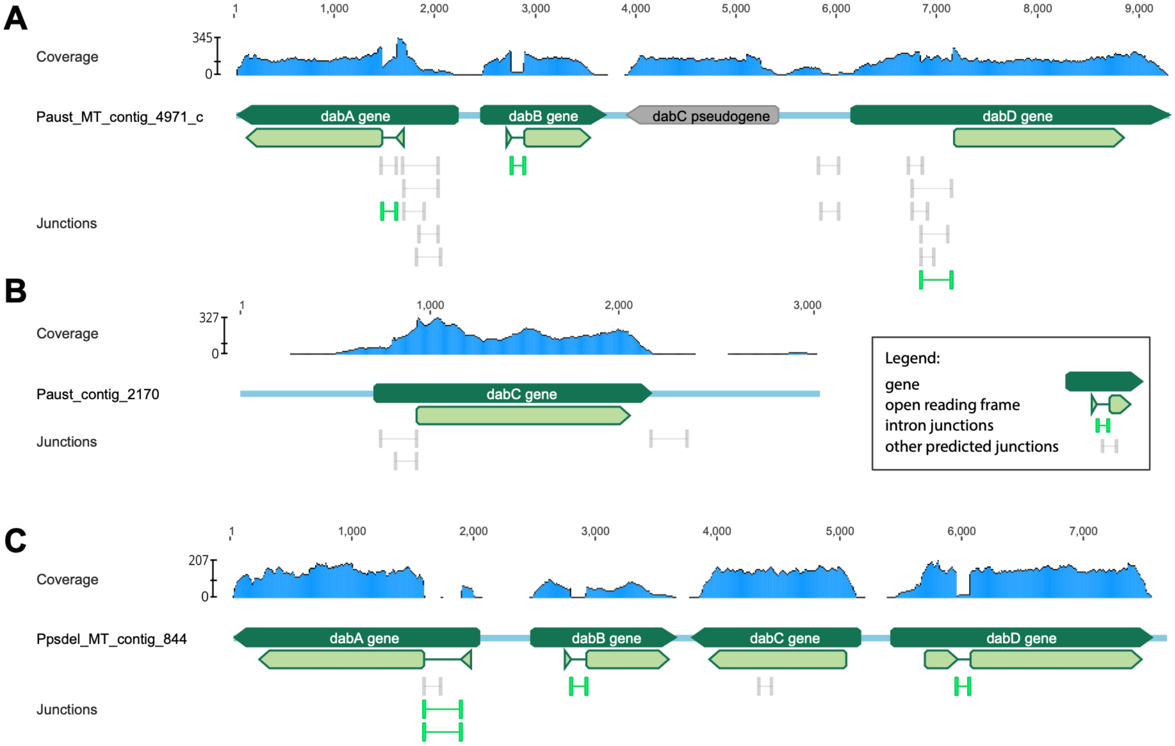


**Fig. S4.** *Dab* gene cluster intron structure based on transcriptomic read mapping. (A) *dab* cluster in *P. australis* MT03; (B) *dabC* in *P. australis* MT03; (C) *dab* cluster in *P.pseudodelicatissima* MT02. Read coverage depth and junctions identified as split reads were compared with gene models manually curated from transcript-to-genome alignments. Data supports the presence of conserved introns in *dabA* and *dabB*, and a novel intron in *P. pseudodelicatissima dabD*. A weakly supported intron appears to be present in the 5'UTR of *P. australis dabD*. Note the spurious expression of the *P. australis dabC* pseudogene, despite the frame-shift mutation introducing a premature stop into the original open reading frame in this locus. Two alternative intron junctions in *P. pseudodelicatissima dabA* reflect allele polymorphism in the splice site. Reads mapped using HISAT2 v2.1.0, visualized using Geneious 2026.1.1. BAM files available in Supplementary Data.

### (A) DabB

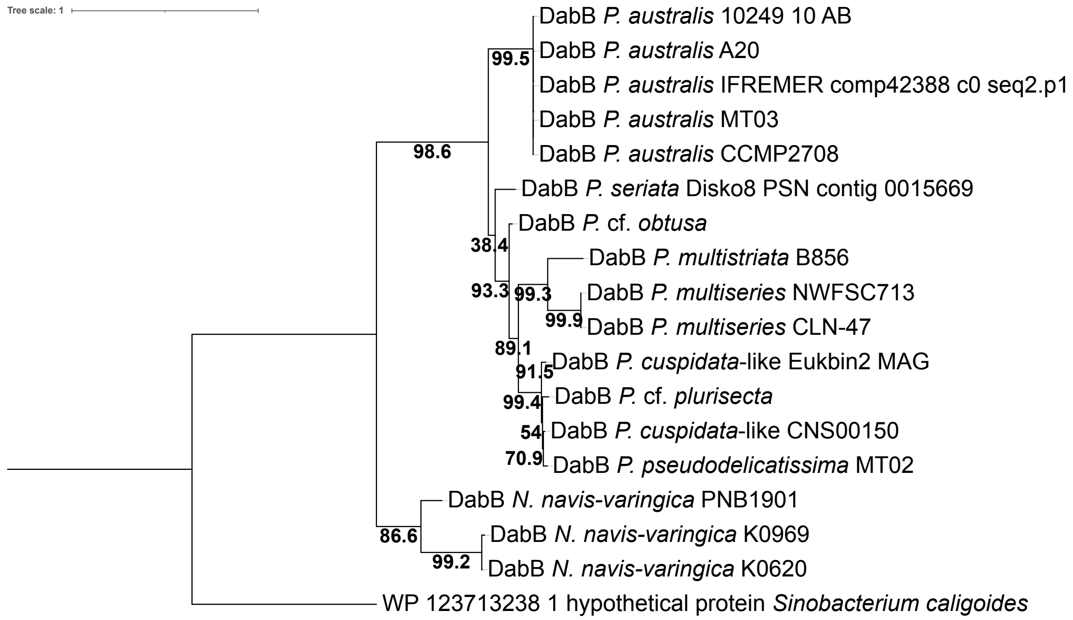


### (B) CYP450 DabD

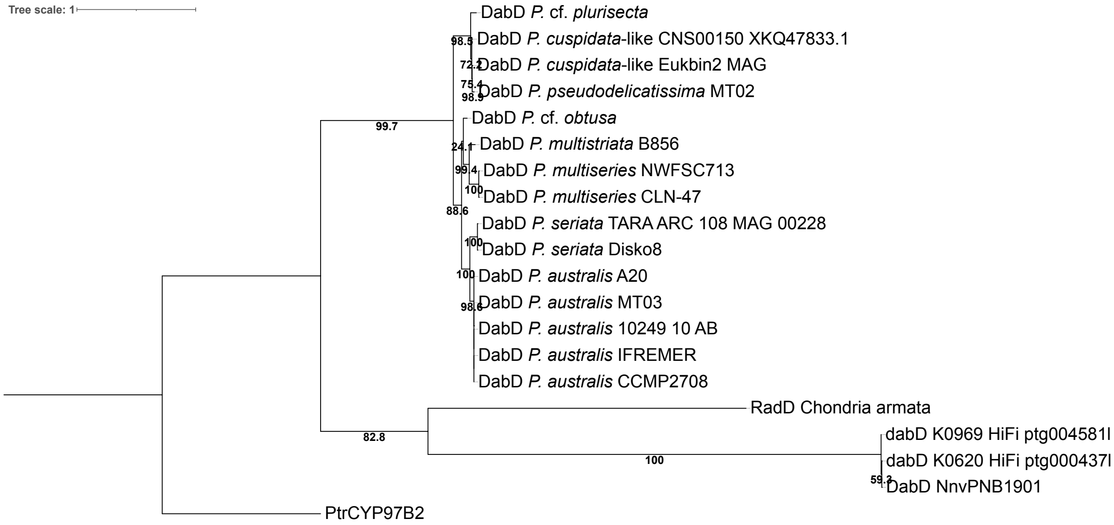


**Fig. S5.** Maximum likelihood phylogenies for single DabB and DabD amino acid sequences with Bootstrap (sh/LRT) shown on branches. Outgroup for DabB was selected based on closest BlastP result. Outgroup for DabD is *Phaeodactylum tricornutum* CYP97B2, which has been biochemically characterized as carotenoid hydroxylase (55). Note different tree scales.

**(A) DabA**

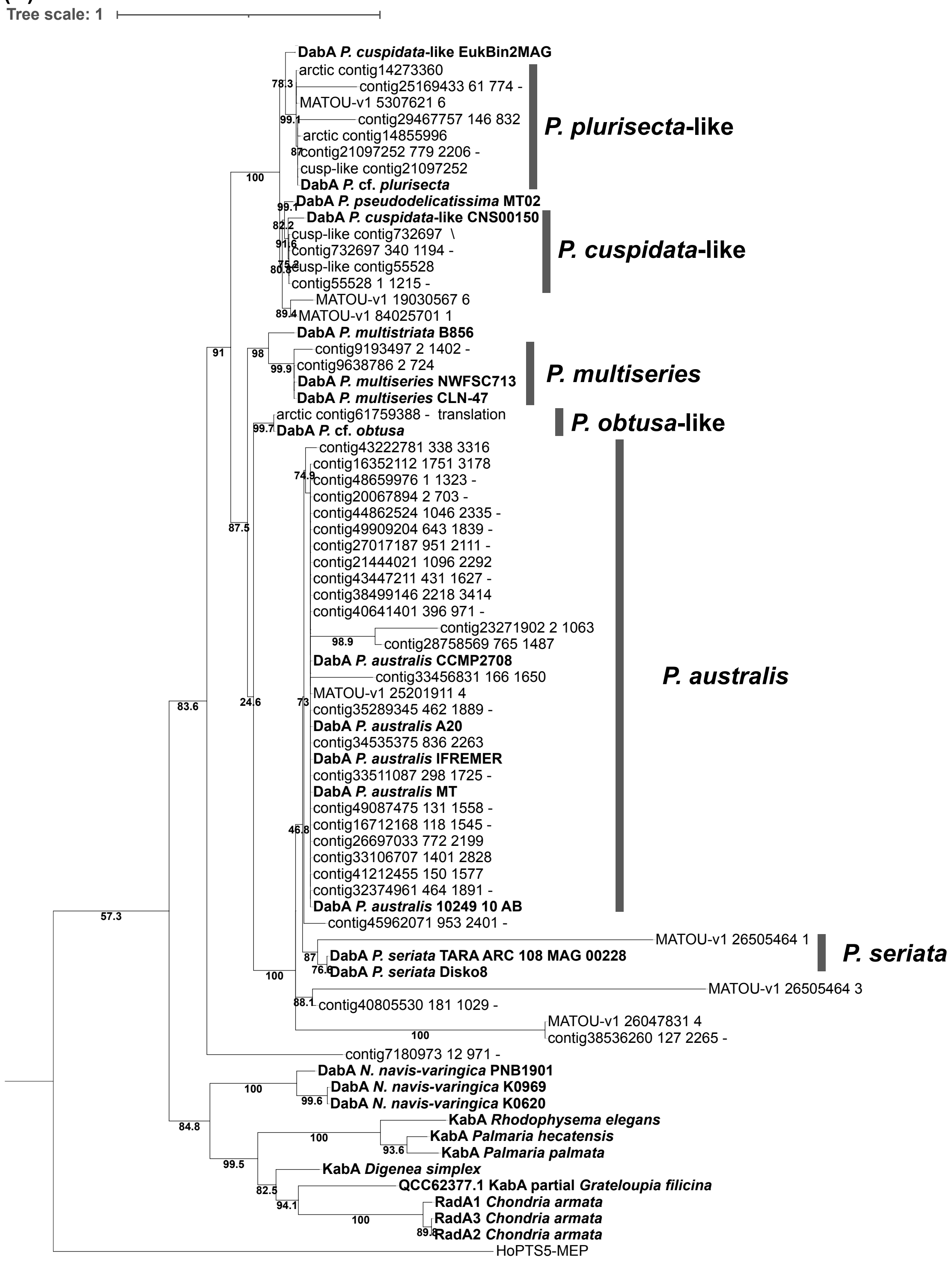


Continued next page

**(B) DabC**

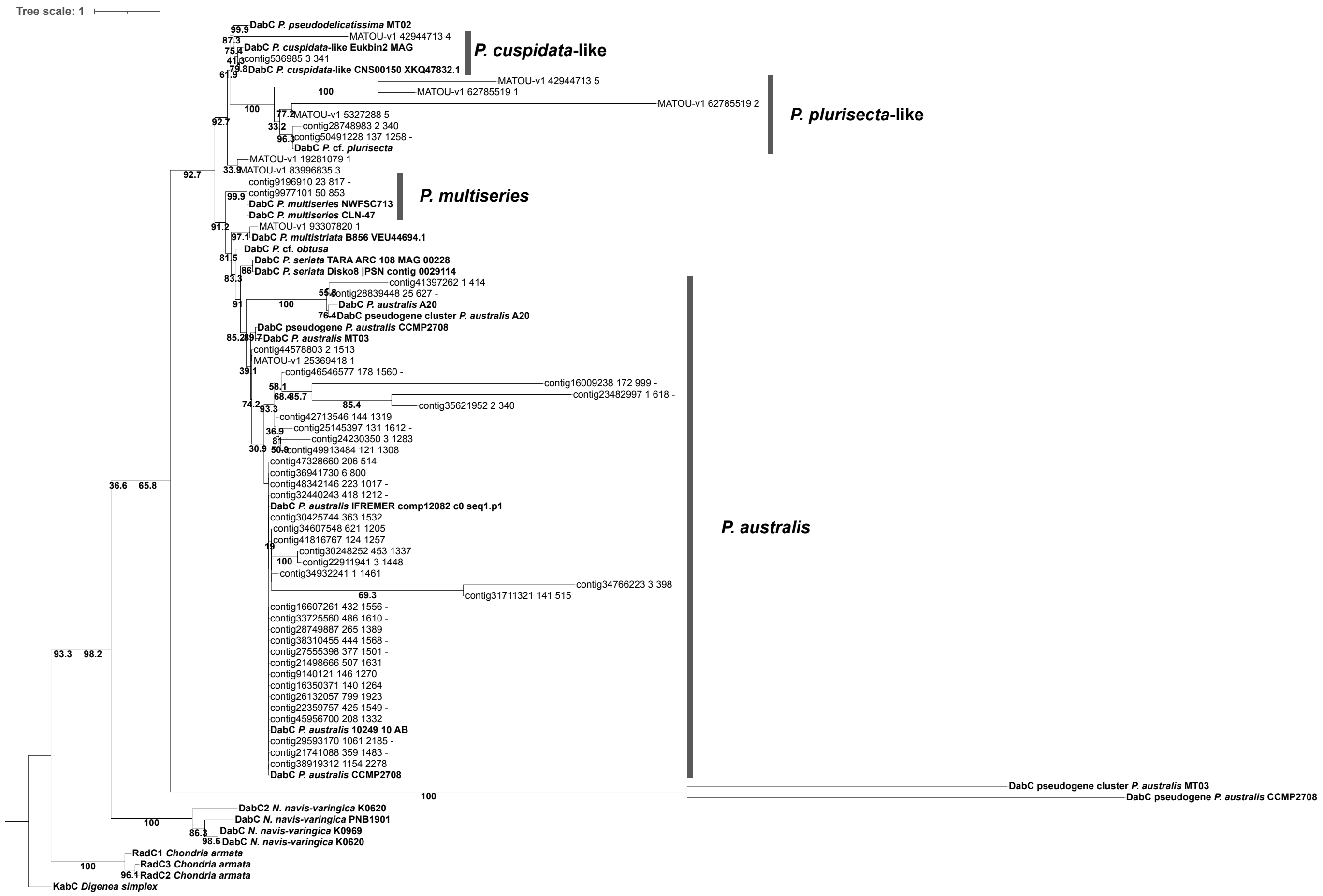


**Fig. S6.** Maximum likelihood phylogenies of environmental transcripts and reference DabA (A) and DabC (B) protein sequences with respective species-level designations. Bootstrap (sh/LRT) values shown on branches. Outgroup for DabA phylogeny is biochemically characterized geranylgeranyl diphosphate synthase HoPTS5 from diatom *Haslea Ostrearia*, likely chloroplast-localized (54), Outgroup for DabC is biochemically characterized kainoid synthase KabC from red alga *Digenea simplex* (56). Note different tree scales.

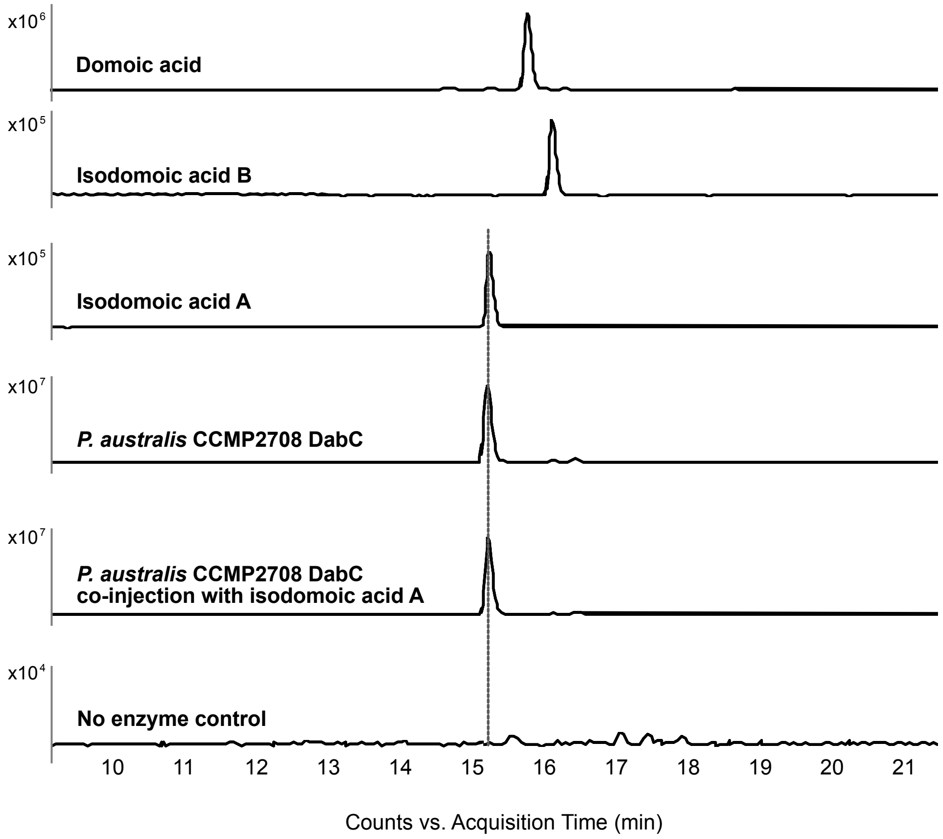


**Fig. S7.** *Pseudo-nitzschia australis* CCMP2708 DabC reaction product with authentic domoic acid and isodomoic acid standards. Negative ion mode LC-MS extracted ion chromatogram (*m*/*z* 310.1±0.1) profiles show, from top to bottom, domoic acid, isodomoic acids C, B, and A, the DabC reaction product, the reaction product co-injected with authentic isodomoic acid A, and a no-enzyme control. The enzymatic product eluted at approximately 15.2 min, matching the isodomoic acid A standard and yielding a single peak upon co-injection, supporting its assignment as isodomoic acid A. No corresponding peak was detected in the no-enzyme control. Signal intensity scales differ among traces, as indicated.

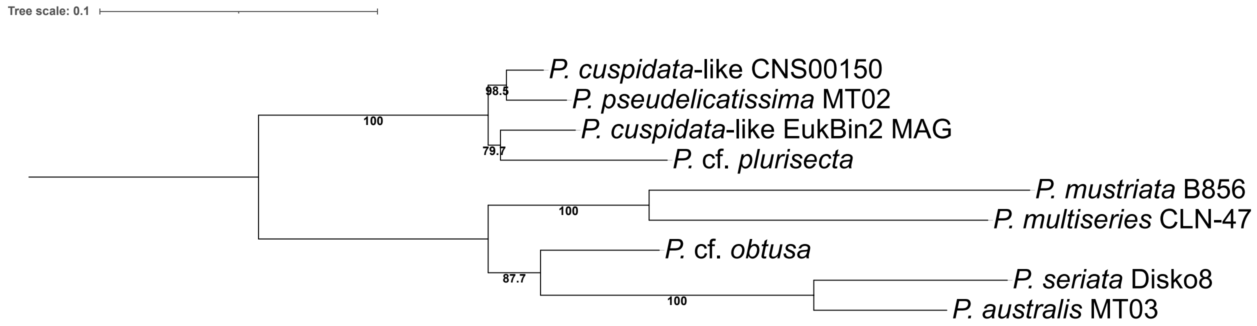


**Fig. S8.** Maximum likelihood phylogeny of concatenated nucleotide sequences of *dab* genes. Bootstrap (sh/LRT) values shown on branches.

**Cell-normalized**

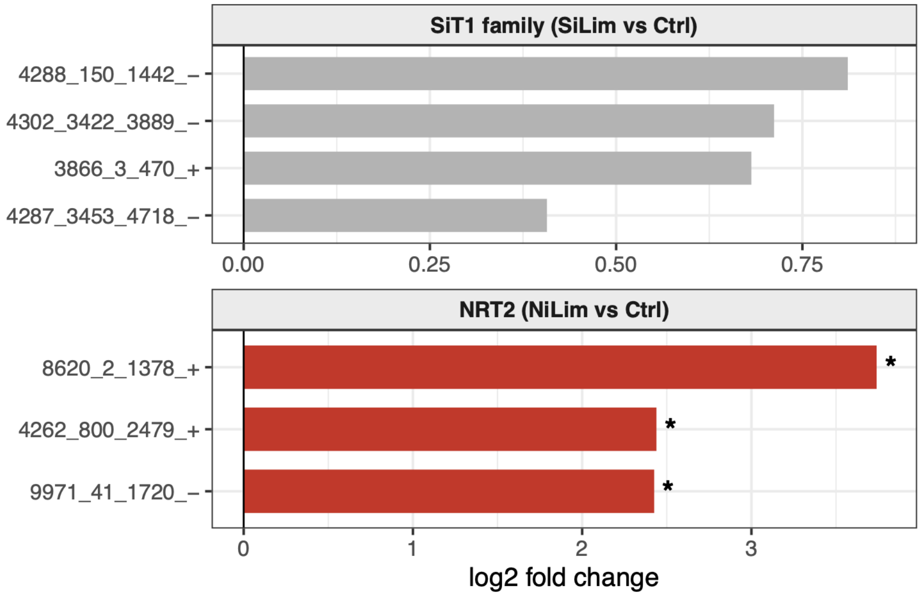


**A. TPM**

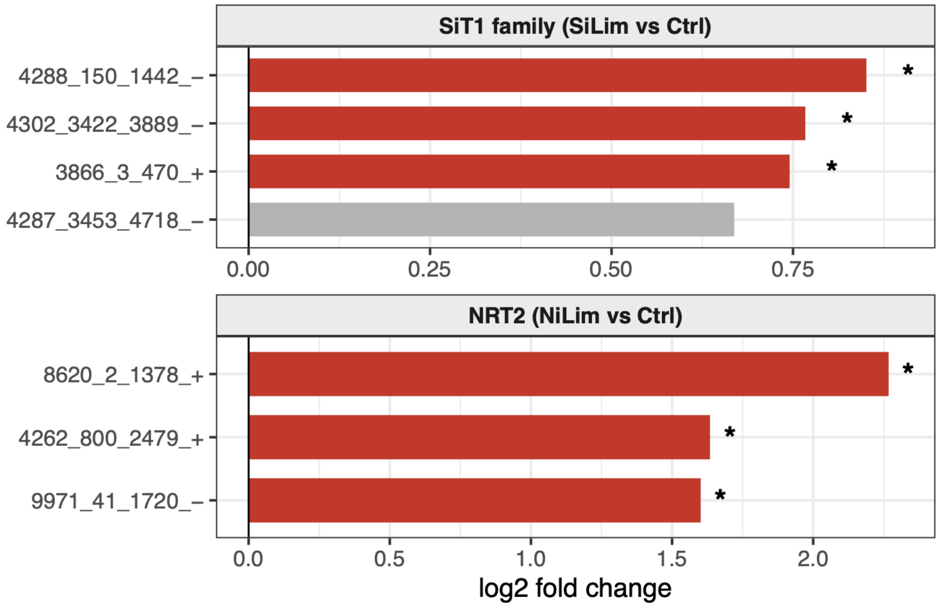


**Fig. S9.** Differential expression of nutrient transporter genes under nitrate and silicate limitation. Log2 fold change values represent the average treatment effect when controlling for time. Results are shown for (A) cell-normalized TMM log2 expression and (B) log2 TPM expression. Bar color (red) and asterisks (*) mark individual genes with significant differential expression (DESeq2, $p < 0.05$). Gene identifiers correspond to contig names from the *P. pseudodelicatissima* metatranscriptome assembly that were confirmed using BlastP searches against reference genes.

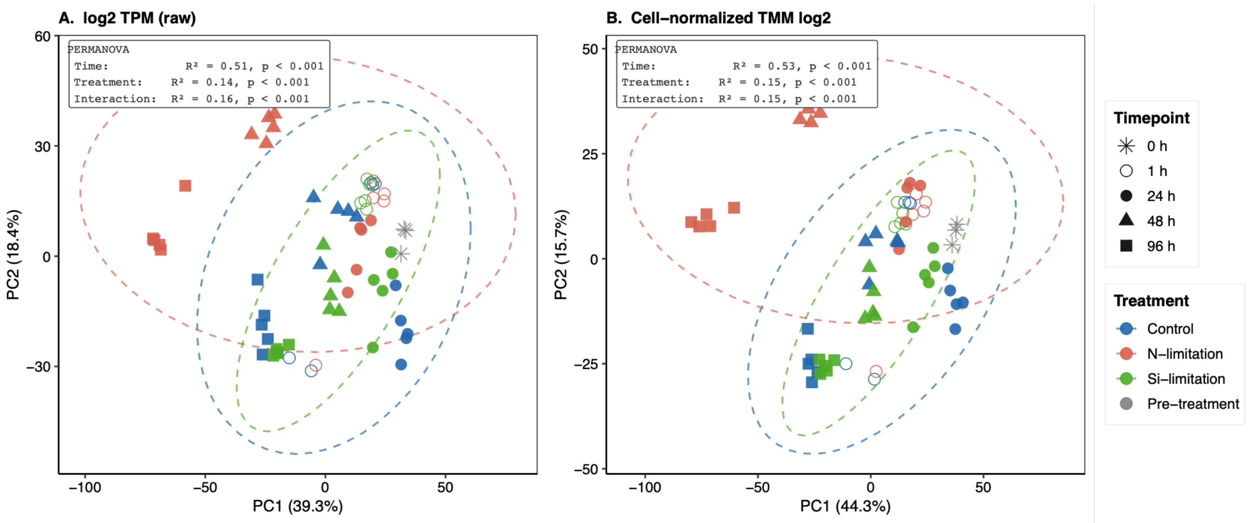


**Fig. S10.** Principal component analysis (PCA) of P. pseudodelicatissima metatranscriptome samples under nutrient limitation. PCA was performed on (A) log2-transformed TPM values and (B) cell-normalized TMM log2 values, using the 2,000 most variable genes across all 63 samples. Points represent individual biological replicates colored by treatment and shaped by timepoint. Dashed ellipses indicate 95% confidence intervals per treatment group. The pre-treatment timepoint (0h; grey asterisks) was included for visualization but excluded from PERMANOVA. PERMANOVA results are shown on each panel; all terms significant at $p < 0.001$ (999 permutations, Euclidean distance). PC1 (39.3% variance in raw TPM; 44.3% in cell-normalized) correlated strongly with time ($r = -0.79$), reflecting progressive transcriptome restructuring over the 96-hour experiment. PC2 (18.4% and 15.7%, respectively) was primarily associated with N-limitation ($r = 0.48$), which drove the largest and most progressive treatment-specific divergence.

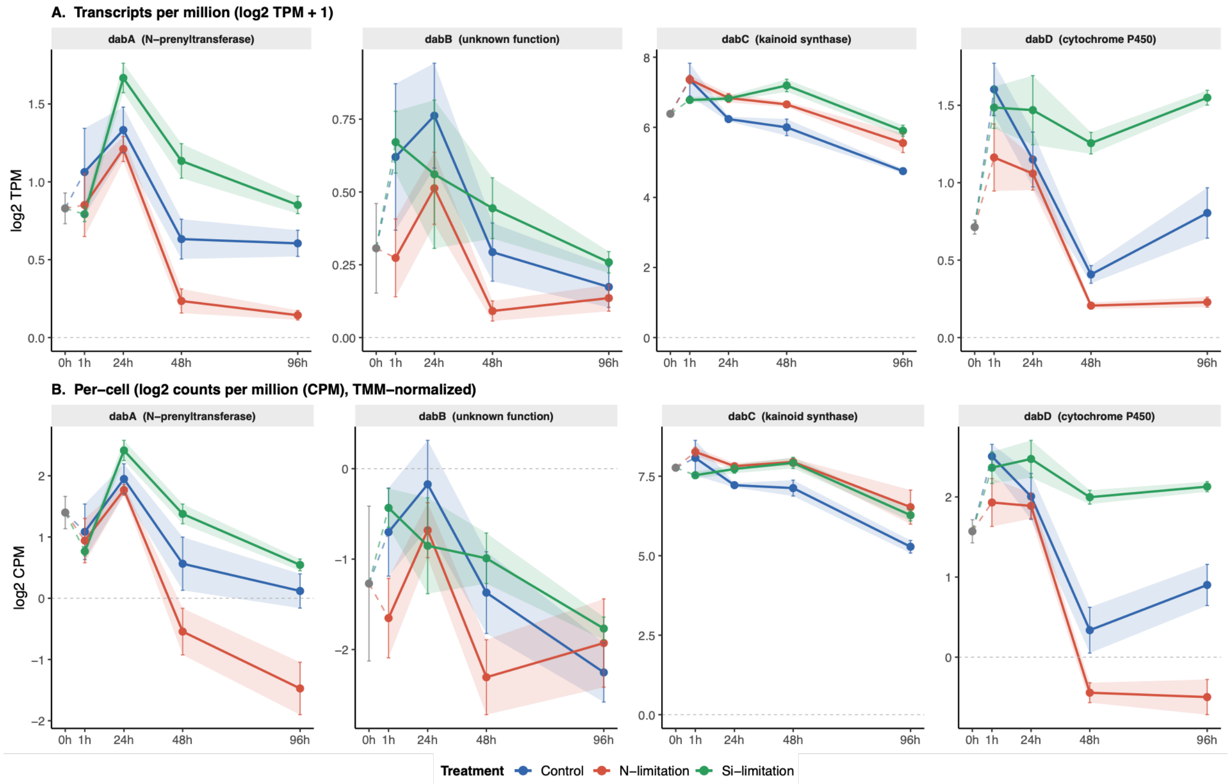


**Fig. S11.** Expression trajectories of domoic acid biosynthesis genes (*dabA–dabD*) across nutrient limitation treatments. (A) Log2-transformed transcripts per million (log2 TPM + 1). (B) Cell-normalized log2 counts per million (log2 CPM), TMM-normalized using edgeR with raw counts scaled to per-cell equivalents prior to library size normalization. Lines represent mean expression ± 1 SE across biological replicates (n = 5 per treatment per timepoint, except the pre-inoculum timepoint Ti = 0 h which is only, n = 3). Shaded ribbons indicate ± 1 standard error for the main time course (1–96 h).

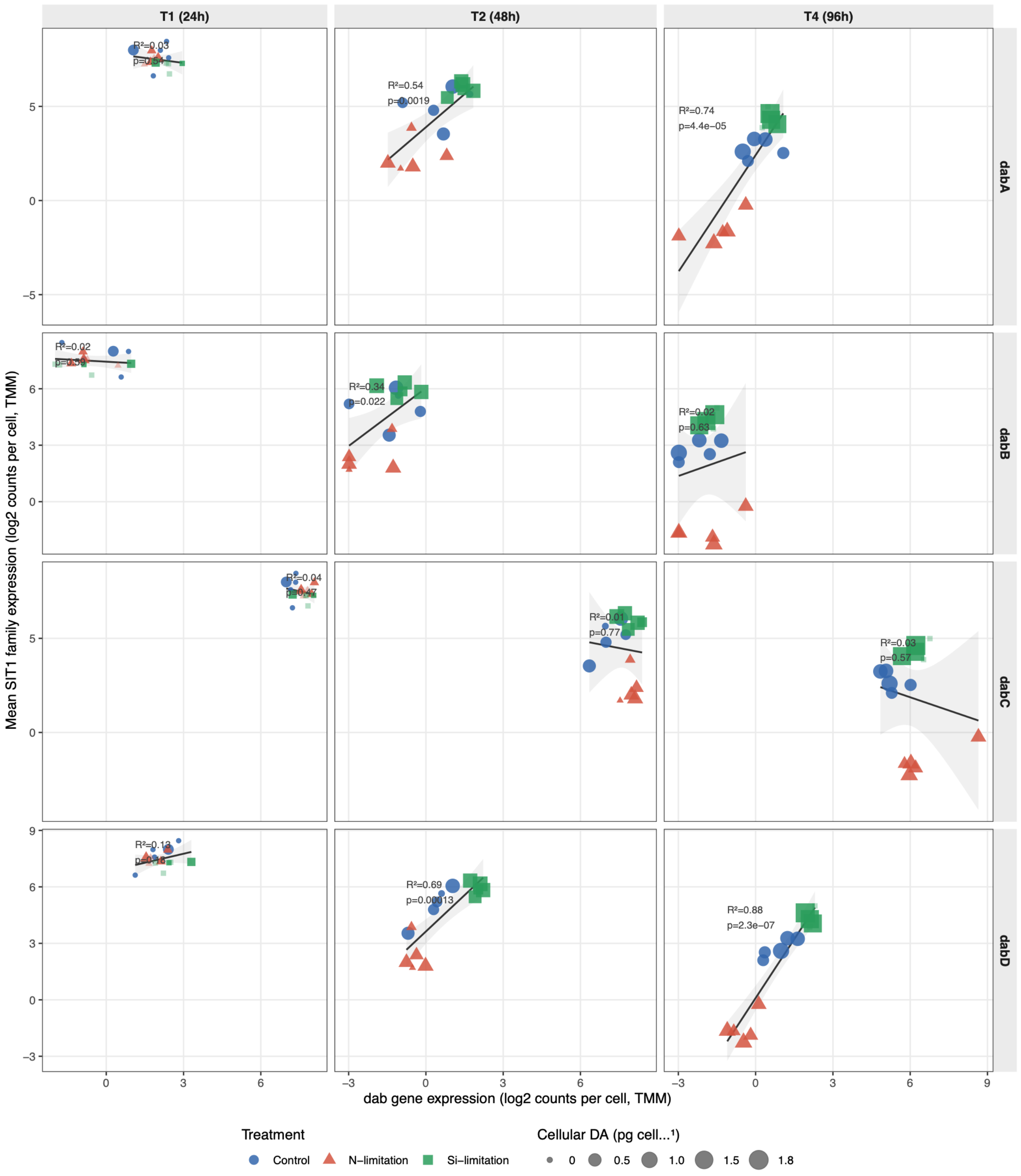


**Fig. S12. Co-expression between individual *dab* genes and the SIT1 silicon transporter family across nutrient limitation treatments.** Points represent individual biological replicates colored by treatment and shaped by treatment identity; point size is proportional to cellular DA concentration (pg cell⁻¹) where available, and faded points indicate samples without DA measurements. The grey line and shaded ribbon show the linear regression fit and 95% confidence interval across all treatments pooled at each timepoint, with $R^2$ and p-values annotated per panel.

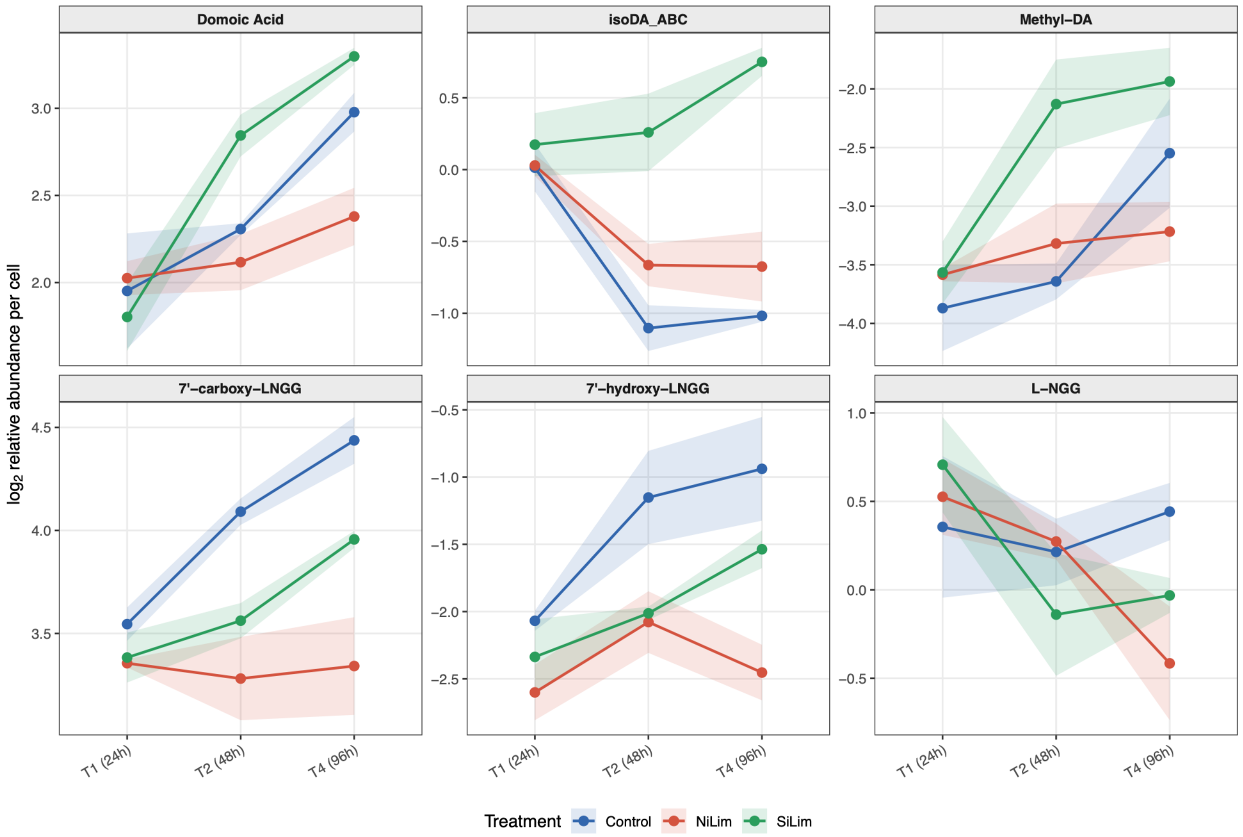


**Fig. S13. Per-cell relative abundance of domoic acid and biosynthetic intermediates across nutrient limitation treatments.** Relative abundance was calculated by transforming PQN-normalized feature intensities to linear scale, dividing by cell density (cells $mL^{-1}$) at each sampling timepoint, and log2-transforming for visualization. Lines represent mean values across biological replicates (n = 3-5 per treatment per timepoint) and ribbons indicate ±1 SE. Feature identities were confirmed by comparison of retention time and MS/MS fragmentation against authentic synthetic standards included in the same LC-MS run, with spectral matching performed through GNPS. Isodomoic acids A, B, and C could not be distinguished by retention time under the LC conditions applied here and are reported as a grouped feature (isoDA_ABC); based on a separate LC analysis of culture extracts, isodomoic acid A is the predominant isomer detected in this strain.

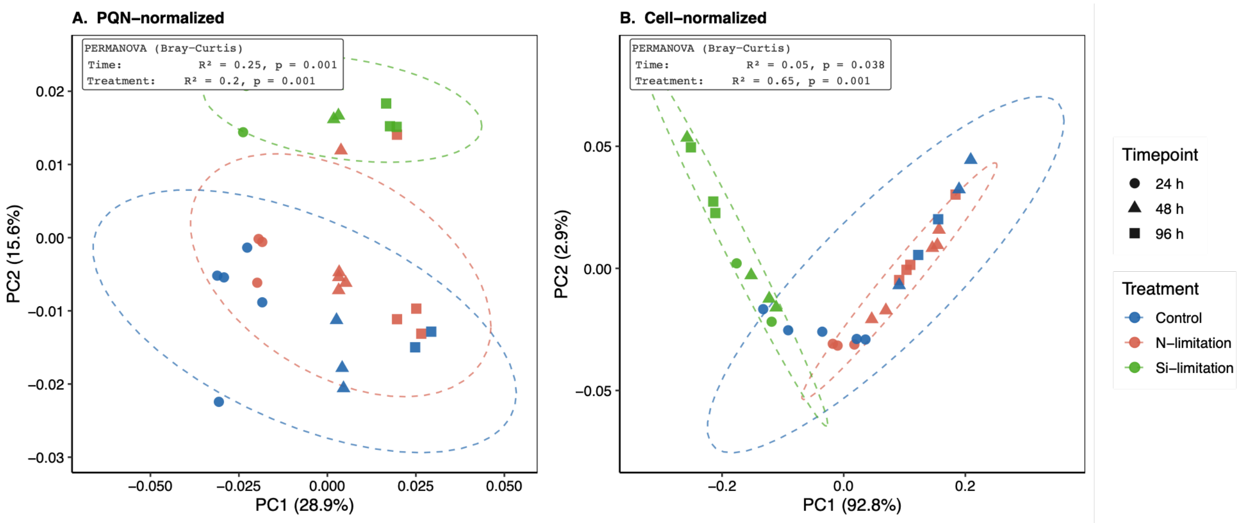


**Fig. S14. Principal coordinates analysis of *P. pseudodelicatissima* holometabolome under nutrient limitation.** Bray-Curtis dissimilarity was calculated on (A) PQN-normalized log2 features and (B) cell-count-normalized log2 feature intensities. Points represent biological replicates. Dashed ellipses indicate 95% confidence intervals per treatment group (t-distribution). PERMANOVA results (Bray-Curtis distance) are annotated. It should be noted that contributions for co-occurring prokaryotes cannot be excluded given culture conditions.

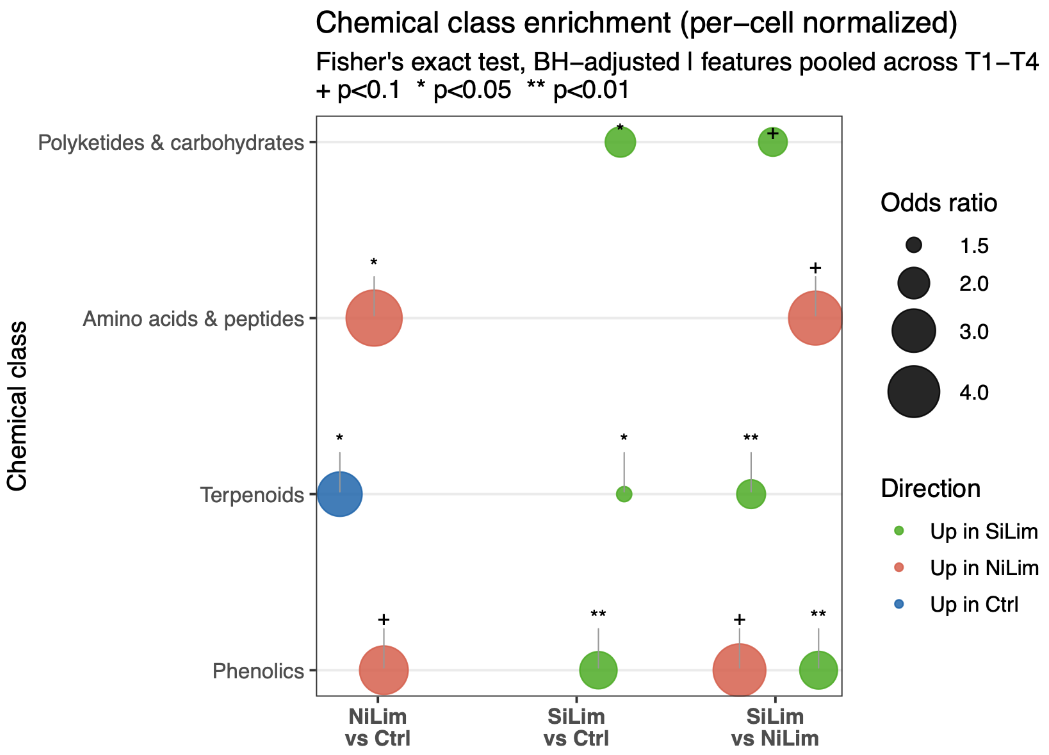


**Fig. S15. Chemical class enrichment of per-cell normalized metabolomics features under nutrient limitation.** Bubble plot showing over-representation of GNPS-generated NPClassifier chemical classes among significantly elevated metabolic features ($p < 0.05$, log2FC > 1) across pairwise treatment comparisons. Features were pooled across timepoints prior to testing. Enrichment was assessed using one-sided Fisher's exact tests comparing the proportion of significantly elevated features in each chemical class against the full tested feature background ($n = 3{,}373$ features). P-values were adjusted for multiple comparisons using the Benjamini-Hochberg method. Bubble size is proportional to the odds ratio, which represents the enrichment of each chemical class among significantly elevated features relative to the full feature background.

**Table S1.** Strain information for those analyzed in this study.

| Strain | Isolation Location | Isolator or Culture Collection |
|---|---|---|
| *Pseudo-nitzschia arenysensis* MT01 | Monterey Wharf, CA, USA | Monica Thukral |
| *Pseudo-nitzschia pseudodelicatissima* MT02 | Monterey Wharf, CA, USA | Monica Thukral |
| *Pseudo-nitzschia australis* MT03 | Monterey Wharf, CA, USA | Monica Thukral |
| *Pseudo-nitzschia hasleana* EBL58 | Monterey Wharf II, CA, USA (36.603683, -121.889275) | Olivia Pawlyk |
| *Pseudo-nitzschia arctica* RCC5316 | Baffin Island, Canada, Arctic Ocean (67.48, -63.78) | Roscoff Culture Collection |
| *Pseudo-nitzschia fraudulenta* CCAP1061/46 | Scottish Association for Marine Science (SAMS) Pontoon, Dunstaffnage, Argyll, Scotland, UK | Culture Collection of Algae and Protozoa (CCAP) |
| *Pseudo-nitzschia pungens* EBL59 | Monterey Wharf II, CA, USA (36.603683, -121.889275) | Olivia Pawlyk, Moss Landing Marine Lab |
| *Pseudo-nitzschia pungens* EBL60 | Monterey Wharf II, CA, USA (36.603683, -121.889275) | Olivia Pawlyk |
| *Pseudo-nitzschia pungens* NWFSC 733-4 | Sequim Bay, WA, USA | Brian Bill |
| *Pseudo-nitzschia australis* A20 | Gulf of Maine, ME, USA | Gloucester Marine Genomics Institute (GMGI) |
| *Pseudo-nitzschia multiseries* NWFSC 713 | Sequim Bay, WA, USA | Brian Bill |
| *Pseudo-nitzschia fraudulenta* F | Gulf of Maine, offshore (44.25, -66.78) | Fish and Wildlife Research Institute (FWC-FWRI) |
| *Pseudo-nitzschia australis* A20 | Gulf of Maine, Bar Harbor (44.434, -68.290) | FWC-FWRI |
| *Pseudo-nitzschia pungens* G20 | Gulf of Maine, Bar Harbor (44.392, -68.204) | FWC-FWRI |

**Table S2.** Summary of the functional annotations for each metatranscriptomics WGCNA module.

| Module | # genes | Mean hub kME | Contains *dab* | Top Pfams |
|---|---|---|---|---|
| turquoise | 1502 | 0.978 | *dabC* | Fasciclin domain; Mycoplasma protein of unknown function, DUF285; Leucine Rich Repeat |
| blue | 1196 | 0.974 | FALSE | DEAD/DEAH box helicase; Helicase conserved C-terminal domain; WD domain, G-beta repeat |
| brown | 1192 | 0.962 | *dabABD* | Chlorophyll A-B binding protein; short chain dehydrogenase; FKBP-type peptidyl-prolyl cis-trans isomerase |
| yellow | 427 | 0.96 | FALSE | PF13715; ABC transporter; TonB-dependent Receptor Plug Domain |
| green | 187 | 0.967 | FALSE | Oxidoreductase family, NAD-binding Rossmann fold; Protein kinase domain; 3-carboxy-cis,cis-muconate lactonizing enzyme |
| red | 113 | 0.963 | FALSE | SOUL heme-binding protein; Mitochondrial matrix Mmp37; Oxidoreductase FAD-binding domain |
| black | 112 | 0.941 | FALSE | Cytochrome b559, alpha (gene psbE) and beta (gene psbF) subunits; ATPase family associated with various cellular activities (AAA); NADH:Ubiquinone/plastoquinone (complex I), various chains |
| pink | 72 | 0.937 | FALSE | Aluminium activated malate transporter; CRAL/TRIO domain; Leucine carboxyl methyltransferase |
| magenta | 57 | 0.934 | FALSE | Bacterial extracellular solute-binding proteins, family 5 Middle; CobW/HypB/UreG, nucleotide-binding domain; Multicopper oxidase |
| purple | 43 | 0.858 | FALSE | Ankyrin repeat; Major Facilitator Superfamily; 7 transmembrane sweet-taste receptor of 3 GCPRs |
| greenyellow | 39 | 0.916 | FALSE | Major Facilitator Superfamily; Na+/H+ antiporter family; Cytidine and deoxycytidylate deaminase zinc-binding region |
| tan | 30 | 0.873 | FALSE | Chlorophyll A-B binding protein; LytB protein; NAD dependent epimerase/dehydratase family |

**Table S3. Differential expression of *dab* biosynthesis genes under nutrient limitation** Per-cell TMM-normalized log2 CPM and TPM log2 values analyzed using R package limma with empirical Bayes moderation. Additive model shows main effects averaged across T1 (24h) – T4 (96h); interaction model identifies timepoint-specific effects. logFC = log2 fold change vs Control; BH-P = Benjamini-Hochberg adjusted p-value.

| Gene | logFC (per-cell) | BH-P (per-cell) | logFC (TPM) | BH-P (TPM) | Significant interaction | BH-P (int.) |
|---|---|---|---|---|---|---|
| **Si-limitation vs Control** | | | | | | |
| ***dabA*** | +0.57 | 0.106 | +0.20 | 0.399 | — | — |
| ***dabB*** | +0.06 | 0.902 | +0.02 | 0.949 | — | — |
| ***dabC*** | +0.77 | **0.012** | +0.59 | **0.021** | — | — |
| ***dabD*** | +1.12 | **0.001** | +0.45 | **0.029** | T2 SiLim (+1.19) | **0.038** |
| **N-limitation vs Control** | | | | | | |
| ***dabA*** | -0.96 | **0.001** | -0.30 | 0.060 | T4 NiLim (-1.41) | **0.044** |
| ***dabB*** | -0.37 | 0.315 | -0.21 | 0.187 | — | — |
| ***dabC*** | +0.89 | **0.001** | +0.52 | **0.012** | — | — |
| ***dabD*** | -0.77 | **0.006** | -0.33 | **0.048** | T4 NiLim (-1.28) | **0.022** |

**Table S4. Significant cross-omics co-expression module correlations.** Pearson correlations between per-cell normalized metatranscriptomic module eigengenes and cell-count-normalized metabolomics module eigengenes, computed across 32 paired samples (BH-adjusted $p < 0.05$). Rows highlighted in blue correspond to correlations with the DA metabolomics module (MEturquoise). Table ordered by BH-adjusted p-value. Relationships with largely unannotated metabolomics modules (n=10) are excluded for simplicity.

| MetaT Module (per-cell) | Metabolomics Module (cell-norm) | MetaT Functional Annotation | Metabolomics Functional Annotation | Pearson r | BH-padj |
|---|---|---|---|---|---|
| brown | green | Chlorophyll/tetrapyrrole biosynthesis, MEP isoprenoid pathway (dabABD) | Membrane metabolism, stress response | +0.652 | 0.0041 |
| brown | blue | Chlorophyll/tetrapyrrole biosynthesis, MEP isoprenoid pathway (dabABD) | Translation, ribosome biogenesis | +0.616 | 0.0056 |
| brown | grey | Chlorophyll/tetrapyrrole biosynthesis, MEP isoprenoid pathway (dabABD) | Unassigned (includes L-NGG) | +0.614 | 0.0056 |
| brown | yellow | Chlorophyll/tetrapyrrole biosynthesis, MEP isoprenoid pathway (dabABD) | Photosynthesis, light harvesting | +0.598 | 0.0070 |
| brown | turquoise | Chlorophyll/tetrapyrrole biosynthesis, MEP isoprenoid pathway (dabABD) | DA module: domoic acid, oxidized LNGG intermediates | +0.580 | 0.0093 |
| brown | black | Chlorophyll/tetrapyrrole biosynthesis, MEP isoprenoid pathway (dabABD) | Lipid, redox metabolism | +0.553 | 0.0138 |
| greenyellow | black | Nitrate/nitrite transport (NRT2), MFS transporters | Lipid, redox metabolism | -0.535 | 0.0138 |
| greenyellow | green | Nitrate/nitrite transport (NRT2), MFS transporters | Membrane metabolism, stress response | -0.538 | 0.0138 |
| greenyellow | grey | Nitrate/nitrite transport (NRT2), MFS transporters | Unassigned (includes L-NGG) | -0.545 | 0.0138 |
| greenyellow | turquoise | Nitrate/nitrite transport (NRT2), MFS transporters | DA module: domoic acid, oxidized LNGG intermediates | -0.549 | 0.0138 |
| greenyellow | yellow | Nitrate/nitrite transport (NRT2), MFS transporters | Photosynthesis, light harvesting | -0.537 | 0.0138 |

| MetaT Module (per-cell) | Metabolomics Module (cell-norm) | MetaT Functional Annotation | Metabolomics Functional Annotation | Pearson r | BH-padj |
|---|---|---|---|---|---|
| greenyellow | blue | Nitrate/nitrite transport (NRT2), MFS transporters | Translation, ribosome biogenesis | -0.523 | 0.0175 |
| green | green | Aromatic catabolism, kinase signaling | Membrane metabolism, stress response | +0.518 | 0.0188 |
| green | grey | Aromatic catabolism, kinase signaling | Unassigned (includes L-NGG) | +0.502 | 0.0240 |
| green | yellow | Aromatic catabolism, kinase signaling | Photosynthesis, light harvesting | +0.498 | 0.0249 |
| green | turquoise | Aromatic catabolism, kinase signaling | DA module: domoic acid, oxidized LNGG intermediates | +0.493 | 0.0254 |
| green | black | Aromatic catabolism, kinase signaling | Lipid, redox metabolism | +0.484 | 0.0274 |
| green | blue | Aromatic catabolism, kinase signaling | Translation, ribosome biogenesis | +0.488 | 0.0274 |
| pink | green | Carbohydrate metabolism, pantothenate biosynthesis | Membrane metabolism, stress response | -0.484 | 0.0274 |
| pink | greenyellow | Carbohydrate metabolism, pantothenate biosynthesis | Lineage-specific regulation | -0.481 | 0.0284 |
| pink | blue | Carbohydrate metabolism, pantothenate biosynthesis | Translation, ribosome biogenesis | -0.476 | 0.0295 |
| pink | grey | Carbohydrate metabolism, pantothenate biosynthesis | Unassigned (includes L-NGG) | -0.476 | 0.0295 |
| red | green | Oxidoreductases, secondary metabolism enzymes | Membrane metabolism, stress response | -0.471 | 0.0316 |
| red | greenyellow | Oxidoreductases, secondary metabolism enzymes | Lineage-specific regulation | -0.464 | 0.0336 |
| pink | turquoise | Carbohydrate metabolism, pantothenate biosynthesis | DA module: domoic acid, oxidized LNGG intermediates | -0.465 | 0.0336 |
| pink | yellow | Carbohydrate metabolism, pantothenate biosynthesis | Photosynthesis, light harvesting | -0.463 | 0.0336 |

| MetaT Module (per-cell) | Metabolomics Module (cell-norm) | MetaT Functional Annotation | Metabolomics Functional Annotation | Pearson r | BH-padj |
|---|---|---|---|---|---|
| pink | black | Carbohydrate metabolism, pantothenate biosynthesis | Lipid, redox metabolism | -0.445 | 0.0429 |
| red | blue | Oxidoreductases, secondary metabolism enzymes | Translation, ribosome biogenesis | -0.445 | 0.0429 |

**Table S5.** Pearson correlations between WGCNA module eigengenes and treatment condition for TPM-normalized and per-cell TMM-normalized metatranscriptomic networks. Delta values indicate the difference in correlation coefficient between per-cell and TPM estimates.

| Module | SiLim TPM | NiLim TPM | SiLim per-cell | NiLim per-cell | SiLim delta | NiLim delta |
|---|---|---|---|---|---|---|
| **green** | 0.25 | -0.18 | 0.47 | -0.93 | 0.23 | -0.75 |
| **brown** | 0.20 | -0.34 | 0.40 | -0.60 | 0.20 | -0.26 |
| **grey** | 0.05 | 0.09 | -0.32 | 0.21 | -0.37 | 0.12 |
| **magenta** | -0.23 | 0.14 | 0.31 | -0.48 | 0.54 | -0.62 |
| **pink** | -0.06 | 0.20 | -0.30 | 0.27 | -0.24 | 0.07 |
| **blue** | -0.20 | 0.59 | -0.26 | 0.78 | -0.06 | 0.18 |
| **red** | 0.08 | -0.02 | -0.16 | 0.07 | -0.24 | 0.10 |
| **yellow** | 0.20 | -0.56 | 0.09 | 0.16 | -0.11 | 0.71 |
| **black** | 0.41 | -0.67 | -0.03 | 0.04 | -0.45 | 0.71 |
| **turquoise** | -0.03 | -0.01 | -0.03 | -0.08 | 0.00 | -0.06 |

**Dataset S1. The *Pseudo-nitzschia* genomic resource sequencing and assembly.** Species, strains, library preparation information, sequencing, and assembly information

**Dataset S2. The *Pseudo-nitzschia* genomic resource annotation.** Species, strains, data sources, annotation counts, assembly, and gene prediction statistics. Source data for Fig. 1.

**Dataset S3. Protein families in *Pseudo-nitzschia* and outgroup diatoms.** Pfam families and OrthoFinder orthogroup distribution. Underlying data for CAFE5.
